# Moderated estimation of fold change and dispersion for RNA-seq data with DESeq2

**DOI:** 10.1101/002832

**Authors:** Michael I Love, Wolfgang Huber, Simon Anders

**Affiliations:** Genome Biology Unit, European Molecular Biology Laboratory, Heidelberg, Germany; Department of Biostatistics and Computational Biology, Dana Farber Cancer Institute and Department of Biostatistics, Harvard School of Public Health,Boston, MA, USA; Department of Computational Molecular Biology, Max Planck Institute for Molecular Genetics, Berlin, Germany

## Abstract

In comparative high-throughput sequencing assays, a fundamental task is the analysis of count data, such as read counts per gene in RNA-seq, for evidence of systematic changes across experimental conditions. Small replicate numbers, discreteness, large dynamic range and the presence of outliers require a suitable statistical approach. We present *DESeq2*, a method for differential analysis of count data, using shrinkage estimation for dispersions and fold changes to improve stability and interpretability of estimates. This enables a more quantitative analysis focused on the strength rather than the mere presence of differential expression. The *DESeq2* package is available at http://www.bioconductor.org/packages/release/bioc/html/DESeq2.html.

## Background

The rapid adoption of high-throughput sequencing (HTS) technologies for genomic studies has resulted in a need for statistical methods to assess quantitative differences between experiments. An important task here is the analysis of RNA-seq data with the aim of finding genes that are differentially expressed across groups of samples. This task is general: methods for it are typically also applicable for other comparative HTS assays, including ChIP-seq, 4C, HiC, or counts of observed taxa in metagenomic studies.

Besides the need to account for the specifics of count data, such as nonNormality and a dependence of the variance on the mean, a core challenge is the small number of samples of typical HTS experiments – often as few as two or three replicates per condition. Inferential methods that treat each gene separately suffer here from lack of power, due to the high uncertainty of within-group variance estimates. In high-throughput assays, this limitation can be overcome by pooling information across genes; specifically, by exploiting assumptions about the similarity of the variances of different genes measured in the same experiment [1].

Many methods for differential expression analysis of RNA-seq data perform such information sharing across genes for variance (or, equivalently, dispersion) estimation. *edgeR* [2,3] moderates the dispersion estimate for each gene toward a common estimate across all genes, or toward a local estimate from genes with similar expression strength, using a weighted conditional likelihood. Our *DESeq* method [4] detects and corrects dispersion estimates which are too low through modeling of the dependence of the dispersion on the average expression strength over all samples. *BBSeq* [5] models the dispersion on the mean, with the mean absolute deviation of dispersion estimates used to reduce the influence of outliers.

*DSS* [6] uses a Bayesian approach to provide an estimate for the dispersion for individual genes which accounts for the heterogeneity of dispersion values for different genes. *baySeq* [7] and *ShrinkBayes* [8] estimate priors for a Bayesian model over all genes, and then provide posterior probabilities or false discovery rates for the case of differential expression.

The most common approach to comparative analysis of transcriptomics data is to test the null hypothesis that the logarithmic fold change (LFC) between treatment and control for a gene’s expression is exactly zero, i.e., that the gene is not at all affected by the treatment. Often the goal of a differential analysis is a list of genes passing multiple-test adjustment, ranked by *p*–value. However, small changes, even if statistically highly significant, might not be the most interesting candidates for further investigation. Ranking by fold-change, on the other hand, is complicated by the noisiness of LFC estimates for genes with low counts. Furthermore, the number of genes called significantly differentially expressed depends as much on the sample size and other aspects of experimental design as it does on the biology of the experiment - and well-powered experiments often generate an overwhelmingly long list of “hits” [9]. We therefore developed a statistical framework to facilitate gene ranking and visualization based on stable estimation of effect sizes (LFCs), as well as testing of differential expression with respect to user-defined thresholds of biological significance.

Here we present *DESeq2*, a successor to our *DESeq* method [4]. *DESeq2* integrates methodological advances with several novel features to facilitate a more quantitative analysis of comparative RNA-seq data by means of shrinkage estimators for dispersion and fold change. We demonstrate the advantages of *DESeq2*’s new features by describing a number of applications possible with shrunken fold changes and their estimates of standard error, including improved gene ranking and visualization, hypothesis tests above and below a threshold, and the “regularized logarithm” transformation for quality assessment and clustering of overdispersed count data. We furthermore compare *DESeq2*’s statistical power with existing tools, revealing that our methodology has high sensitivity and precision, while controlling the false positive rate. *DESeq2* is available as an R/Bioconductor package [10] at http://www.bioconductor.org/packages/release/bioc/html/DESeq2.html.

## Results and discussion

### Model and normalization

The starting point of a *DESeq2* analysis is a count matrix *K* with one row for each gene *i* and one column for each sample *j*. The matrix entries *K_ij_* indicate the number of sequencing reads that have been unambiguously mapped to a gene in a sample. Note that although we refer in this paper to counts of reads in genes, the methods presented here can be applied as well to other kinds of HTS count data. For each gene, we fit a generalized linear model (GLM) [11] as follows.

We model read counts *K_ij_* as following a Negative Binomial distribution (sometimes also called a Gamma-Poisson distribution) with mean *μ_ij_* and dispersion *α_i_*. The mean is taken as a quantity *q_ij_*, proportional to the concentration of cDNA fragments from the gene in the sample, scaled by a normalization factor *S_ij_*, i.e., *μ_ij_* = *s_ij_q_ij_*. For many applications, the same constant *s_j_* can be used for all genes in a sample, which then accounts for differences in sequencing depth between samples. To estimate these *size factors*, the *DESeq2* package offers the median-of-ratios method already used in *DESeq* [4]. However, it can be advantageous to calculate gene-specific normalization factors *s_ij_* to account for further sources of technical biases such as differing dependence on GC content, gene length or the like, using published methods [12,13], and these can be supplied instead.

We use GLMs with logarithmic link, log_2_ *q_ij_* = ∑_*r*_ *x_jr_β_ir_*, with design matrix elements *x_jr_* and coefficients *β_ir_*. In the simplest case of a comparison between two groups, such as treated and control samples, the design matrix elements indicate whether a sample *j* is treated or not, and the GLM fit returns coefficients indicating the overall expression strength of the gene and the log_2_ fold change between treatment and control. The use of linear models, however, provides the flexibility to also analyze more complex designs, as is often useful in genomic studies [14].

**Figure 1.**
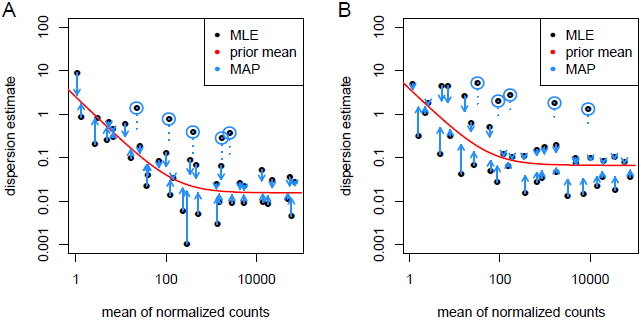
Shrinkage estimation of dispersion. Plot of dispersion estimates over the average expression strength (A) for the Bottomly et al. [15] dataset with 6 samples across 2 groups and (B) for 5 samples from the Pickrell et al. [16] dataset, fitting only an intercept term. First, gene-wise maximum likelihood estimates (MLE) are obtained using only the respective gene’s data (black dots). Then, a curve (red) is fit to the MLEs to capture the overall trend of dispersion-mean dependence. This fit is used as a prior mean for a second estimation round, which results in the final maximum *a posteriori* (MAP) estimates of dispersion (arrow heads). This can be understood as a shrinkage (along the blue arrows) of the noisy gene-wise estimates toward the consensus represented by the red line. The black points circled in blue are detected as dispersion outliers and not shrunk toward the prior (shrinkage would follow the dotted line). For clarity, only a subset of genes is shown, which is enriched for dispersion outliers. Supplementary Figure S1 displays the same data but with dispersions of all genes shown.

### Empirical Bayes shrinkage for dispersion estimation

Within-group variability, i.e., the variability between replicates, is modeled by the dispersion parameter *α_i_*, which describes the variance of counts via Var 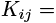 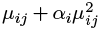. Accurate estimation of the dispersion parameter *α_i_* is critical for the statistical inference of differential expression. For studies with large sample sizes this is usually not a problem. For controlled experiments, however, sample sizes tend to be smaller (experimental designs with as little as two or three replicates are common and reasonable), resulting in highly variable dispersion estimates for each gene. If used directly, these noisy estimates would compromise the accuracy of differential expression testing.

One sensible solution is to share information across genes. In *DESeq2*, we assume that genes of similar average expression strength have similar dispersion. We here explain the concepts of our approach using as example a dataset by Bottomly et al. [15] with RNA-seq data of mice of two different strains and a dataset by Pickrell et al. [16] with RNA-seq data of human lymphoblastoid cell lines. For the mathematical details, see Methods.

We first treat each gene separately and estimate “gene-wise” dispersion estimates (using maximum likelihood), which rely only on the data of each individual gene (black dots in Figure 1). Next, we determine the location parameter of the distribution of these estimates; to allow for dependence on average expression strength, we fit a smooth curve, as shown by the red line in Figure 1. This provides an accurate estimate for the expected dispersion value for genes of a given expression strength but does not represent deviations of individual genes from this overall trend. We then shrink the gene-wise dispersion estimates toward the values predicted by the curve to obtain final dispersion values (blue arrow heads). We use an empirical Bayes approach (Methods), which lets the strength of shrinkage depend (i) on an estimate of how close true dispersion values tend to be to the fit and (ii) on the degrees of freedom: as the sample size increases, the shrinkage decreases in strength, and eventually becomes negligible. Our approach therefore accounts for gene-specific variation to the extent that the data provide this information, while the fitted curve aids estimation and testing in less information-rich settings.

Our approach is similar to the one used by *DSS* [6], in that both methods sequentially estimate a prior distribution for the true dispersion values around the fit, and then provide the maximum *a posteriori* as the final estimate. It differs from the previous implementation of *DESeq*, which used the maximum of the fitted curve and the gene-wise dispersion estimate as the final estimate and tended to overestimate the dispersions (Supplementary Figure S2). The approach of *DESeq2* differs from that of *edgeR* [3], as *DESeq2* estimates the width of the prior distribution from the data and therefore automatically controls the amount of shrinkage based on the observed properties of the data. In contrast, the default steps in *edgeR* require a user-adjustable parameter, the *prior degrees of freedom*, which weighs the contribution of the individual gene estimate and *edgeR’s* dispersion fit.

Note that in Figure 1 a number of genes with gene-wise dispersion estimates below the curve have their final estimates raised substantially. The shrinkage procedure thereby helps avoid potential false positives which can result from underestimates of dispersion. If, on the other hand, an individual gene’s dispersion is far above the distribution of the gene-wise dispersion estimates of other genes, then the shrinkage would lead to a greatly reduced final estimate of dispersion. We reasoned that in many cases, the reason for extraordinarily high dispersion of a gene is that it does not obey our modeling assumptions; some genes may show much higher variability than others for biological or technical reasons, even though they have the same average expression levels. In these cases, inference based on the shrunken dispersion estimates could lead to undesirable false positive calls. *DESeq2* handles these cases by using the gene-wise estimate instead of the shrunken estimate when the former is more than 2 residual standard deviations above the curve.

### Empirical Bayes shrinkage for fold-change estimation

A common difficulty in the analysis of HTS data is the strong variance of logarithmic fold change estimates (LFCs) for genes with low read count. We demonstrate this issue using the dataset by Bottomly et al. [15]. As visualized in Figure 2A, weakly expressed genes seem to show much stronger differences between the compared mouse strains than strongly expressed genes. This phenomenon, seen in most HTS datasets, is a direct consequence of the fact that one is dealing with *count* data, in which ratios are inherently noisier when counts are low. This heteroskedasticity (variance of LFCs depending on mean count) complicates downstream analysis and data interpretation, as it makes effect sizes difficult to compare across the dynamic range of the data.

*DESeq2* overcomes this issue by shrinking LFC estimates toward zero in a manner such that shrinkage is stronger when the available information for a gene is low, which may be because counts are low, dispersion is high, or there are few degrees of freedom. We again employ an empirical Bayes procedure: we first perform ordinary GLM fits to obtain maximum-likelihood estimates (MLE) for the LFCs and then fit a zero-centered Normal distribution to the observed distribution of MLEs over all genes. This distribution is used as a prior on LFCs in a second round of GLM fits, and the maximum *a posteriori* (MAP) estimates are kept as final estimates of LFC. Furthermore, a standard error for each estimate is reported, which is derived from the posterior’s curvature at its maximum (see Methods for details). These shrunken LFCs and their standard errors are used in the Wald tests for differential expression described in the next section.

**Figure 2.**
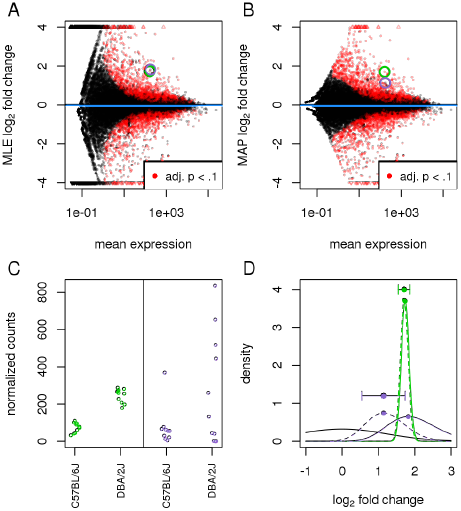
Effect of shrinkage on logarithmic fold change estimates. Plots of the (A) maximum likelihood estimate (MLE, i.e., no shrinkage) and (B) maximum *a posteriori* (MAP) estimate (i. e., with shrinkage) for the logarithmic fold changes attributable to mouse strain, over the average expression strength for a 10 vs 11 sample comparison of the Bottomly et al. [15] dataset. Small triangles at the top and bottom of the plots indicate points that would fall outside of the plotting window. Two genes with similar mean count and MLE logarithmic fold change are highlighted with green and purple circles. (C) The counts (normalized by size factors *S_j_*) for these genes reveal low dispersion for the gene in green and high dispersion for the gene in purple. (D) Density plots of the likelihoods (solid lines, scaled to integrate to 1) and the posteriors (dashed lines) for the green and purple gene and of the prior (solid black line): due to the higher dispersion of the purple gene, its likelihood is wider and less peaked (indicating less information), and the prior has more influence on its posterior than in the case of the green gene. The stronger curvature of the green posterior at its maximum translates to a smaller reported standard error for the MAP LFC estimate (horizontal error bar).

The resulting MAP LFCs are biased toward zero in a manner that removes the problem of “exaggerated” LFCs for low counts. As Figure 2B shows, the strongest LFCs are no longer exhibited by genes with weakest expression. Rather, the estimates are more evenly spread around zero, and for very weakly expressed genes (less than one read per sample on average), LFCs hardly deviate from zero, reflecting that accurate LFC estimates are not possible here.

The strength of shrinkage does not depend simply on the mean count, but rather on the amount of information available for the fold change estimation (as indicated by the observed Fisher information; see Methods). Two genes with equal expression strength but different dispersions will experience different amount of shrinkage (Figure 2C–D). The shrinkage of LFC estimates can be described as a “bias-variance trade-off” [17]: for genes with little information for LFC estimation, a reduction of the strong variance is “bought” at the cost of accepting a bias toward zero, and this can result in an overall reduction in mean squared error, e. g., when comparing to LFC estimates from a new dataset. Genes with high information for LFC estimation will have, in our approach, LFCs with both low bias and low variance. Furthermore, as the degrees of freedom increase, and the experiment provides more information for LFC estimation, the shrunken estimates will converge to the unshrunken estimates. We note that other Bayesian efforts toward moderating fold changes for RNA-seq include hierarchical models [8,18] and the *GFOLD* (or “generalized fold change”) tool [19], which uses a posterior distribution of logarithmic fold changes.

**Figure 3.**
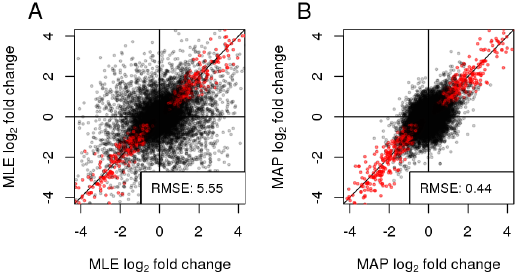
Stability of logarithmic fold changes. *DESeq2* is run on equally split halves of the data of Bottomly et al. [15], and the logarithmic fold changes from the halves are plotted against each other, (A) showing MLEs, i.e., without LFC shrinkage, (B) showing MAP estimates, i.e., with shrinkage. Points in the top left and bottom right quadrant indicate genes with a change of sign of logarithmic fold change. Red points indicate genes with adjusted *p*–value less than 0.1. The legend displays the root mean squared error of the estimates in group I to those in group II.

The shrunken MAP LFCs offer a more reproducible quantification of transcriptional differences than standard MLE LFCs. To demonstrate this, we split the Bottomly et al. samples equally into two groups, I and II, such that each group contained a balanced split of the strains, simulating a scenario where an experiment (samples in group I) is performed, analyzed and reported, and then independently replicated (samples in group II). Within each group, we estimated LFCs between the strains and compared between group I and II, using the MLE LFCs (Figure 3A) and using the MAP LFCs (Figure 3B). Because the shrinkage moves large LFCs that are not well supported by the data toward zero, the agreement between the two independent sample groups increases considerably. Therefore, shrunken fold-change estimates offer a more reliable basis for quantitative conclusions than normal maximum-likelihood estimates.

This makes shrunken LFCs also suitable for ranking genes, e. g., to prioritize them for follow-up experiments. For example, if we sort the genes in the two sample groups of Figure 3 by unshrunken LFC estimates, and consider the 100 genes with the strongest up-or down-regulation in group I, we find only 21 of these again among the top 100 up-or down-regulated genes in group II. However, if we rank the genes by shrunken LFC estimates, the overlap improves to 81 of 100 genes (Supplementary Figure S3).

A simpler, often used method is to add a fixed number (“pseudocount”) to all counts before forming ratios. However, this requires the choice of a tuning parameter and only reacts to one of the sources of uncertainty, low counts, but not to gene-specific dispersion differences, or sample size. We demonstrate this in the *Benchmarks* section below.

**Figure 4.**
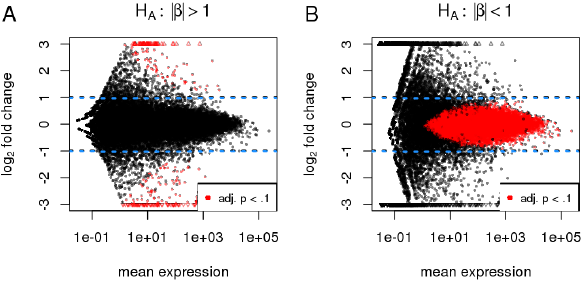
Hypothesis testing involving non-zero thresholds. Shown are MA-plots for a 10 vs 11 comparison using the Bottomly et al. [15] dataset, with highlighted points indicating low adjusted *p*–values. The alternate hypotheses are that logarithmic (base 2) fold changes are (A) greater than 1 in absolute value or (B) less than 1 in absolute value.

### Hypothesis tests for differential expression

After GLMs are fit for each gene, one may test for each model coefficient whether it differs significantly from zero. To this end, *DESeq2* reports the standard error for each shrunken LFC estimate, obtained from the curvature of the coefficient’s posterior (dashed lines in Figure 2D) at its maximum. For significance testing, *DESeq2* uses a Wald test: the shrunken estimate of LFC is divided by its standard error, resulting in a *z*–statistic which is compared to a standard Normal. (See Methods for details.) The Wald test allows testing of individual coefficients, or contrasts of coefficients, without the need to fit a reduced model as with the likelihood ratio test, though the likelihood ratio test is also available as an option in *DESeq2*. The Wald test *p*–values from the subset of genes that pass an independent filtering step, described in the next section, are adjusted for multiple testing using the procedure of Benjamini and Hochberg [20].

### Automatic independent filtering

Due to the large number of tests performed in the analysis of RNA-seq and other genome-wide experiments, the multiple testing problem needs to be addressed. A popular objective is control or estimation of the false discovery rate (FDR). Multiple testing adjustment tends to be associated with a loss of power, in the sense that the false discovery rate for a set of genes is often higher than the individual *p*–values of these genes. However, the loss can be reduced if genes are omitted from the testing that have little or no chance of being detected as differentially expressed, provided that the criterion for omission is independent of the test statistic under the null hypothesis [21] (see Methods). *DESeq2* uses the average expression strength of each gene, across all samples, as its filter criterion, and it omits all genes with mean normalized counts below a filtering threshold from multiple testing adjustment. *DESeq2* by default will choose a threshold that maximizes the number of genes found at a user-specified target FDR. In Figure 2A–Figure 2B and Figure 3, genes found in this way to be significant at an estimated FDR of 10% are depicted in red.

Depending on the distribution of mean normalized counts, the resulting increase in power can be substantial, sometimes making the difference whether or not any differentially expressed genes are detected.

## Hypothesis tests with thresholds on effect size

### Specifying minimum effect size

Most approaches to testing for differential expression, including the default approach of *DESeq2*, test against the null hypothesis of *zero* logarithmic fold change. However, once any biological processes are genuinely affected by the difference in experimental treatment, this null hypothesis implies that the gene under consideration is *perfectly* decoupled from these processes. Due to the high interconnectedness of cells’ regulatory networks, this hypothesis is, in fact, implausible, and arguably wrong for many if not most genes. Consequently, with sufficient sample size, even genes with a very small, but non-zero logarithmic fold change will eventually be detected as differentially expressed. A change should therefore be of sufficient magnitude to be considered *biologically significant*. For small scale experiments, statistical significance is often a much stricter requirement than biological significance, thereby relieving the researcher from the need to decide on a threshold for biological significance.

For well-powered experiments, however, a statistical test against the conventional null hypothesis of zero logarithmic fold change may report genes with statistically significant changes that are so weak in effect strength that they could be considered irrelevant or distracting. A common procedure is to disregard genes whose estimated logarithmic fold change *β_ir_* is below some threshold, |*β_ir_*| ≤ *θ*. However, this approach loses the benefit of an easily interpretable false discovery rate, as the reported *p*–value and adjusted *p*–value still correspond to the test of *zero* logarithmic fold change. It is therefore desirable to include the threshold into the statistical testing procedure directly, i. e., not to filter post-hoc on a reported fold-change *estimate*, but rather to statistically evaluate directly whether there is sufficient evidence that the logarithmic fold change is above the chosen threshold.

*DESeq2* offers tests for composite null hypotheses of the form |*β_ir_*|≤ θ, where *β_ir_* is the shrunken LFC from the estimation procedure described above. (See Methods for details.) Figure 4A demonstrates how such a thresholded test gives rise to a curved decision boundary: to reach significance, the estimated LFC has to exceed the specified threshold by an amount that depends on the available information. We note that related approaches to generate gene lists that satisfy both statistical and biological significance criteria have been previously discussed for microarray data [22] and recently for sequencing data [18].

### Specifying maximum effect size

Sometimes, a researcher is interested in finding genes that are not, or only very weakly, affected by the treatment or experimental condition. This amounts to a setting similar to the one just discussed, but the roles of null and alternative hypotheses are swapped. We are here asking for evidence of the effect being weak, not for evidence of the effect being zero, because the latter question is rarely tractable. The meaning of *weak* needs to be quantified for the biological question at hand by choosing a suitable threshold θ for the LFC. For such analyses, *DESeq2* offers a test of the composite null hypothesis |*β_ir_*|≥ θ, which will report genes as significant for which there is evidence that their LFC is weaker than θ. Figure 4B shows the outcome of such a test. For genes with very low read count, even an estimate of zero LFC is not significant, as the large uncertainty of the estimate does not allow us to exclude that the gene may in truth be more than weakly affected by the experimental condition. Note the lack of LFC shrinkage: To find genes with weak differential expression, *DESeq2* requires that the LFC shrinkage has been disabled. This is because the zero-centered prior used for LFC shrinkage embodies a *prior* belief that LFCs tend to be small, and hence is inappropriate here.

### Detection of count outliers

Parametric methods for detecting differential expression can have gene-wise estimates of logarithmic fold change overly influenced by individual outliers that do not fit the distributional assumptions of the model [23]. An example of such an outlier would be a gene with single-digit counts for all samples, except one sample with a count in the thousands. As the aim of differential expression analysis is typically to find *consistently* up-or down-regulated genes, it is useful to consider diagnostics for detecting individual observations which overly influence the logarithmic fold change estimate and *p*–value for a gene. A standard outlier diagnostic is Cook’s distance [24], which is defined within each gene for each sample as the scaled distance that the coefficient vector, 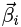, of a linear or generalized linear model would move if the sample were removed and the model refit.

*DESeq2* flags, for each gene, those samples which have a Cook’s distance greater than the 0.99 quantile of the *F(p,m – p)* distribution, where *p* is the number of model parameters including the intercept, and *m* is the number of samples. The use of the *F* distribution is motivated by the heuristic reasoning that removing a single sample should not move the vector 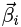 outside of a 99% confidence region around 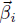 fit using all the samples [24]. However, if there are 2 or fewer replicates for a condition, these samples do not contribute to outlier detection, as there are insufficient replicates to determine outlier status.

How should one deal with flagged outliers? In an experiment with many replicates, discarding the outlier and proceeding with the remaining data might make best use of the available data. In a small experiment with few samples, however, the presence of an outlier can impair inference regarding the affected gene, and merely ignoring the outlier may even be considered data cherry-picking – and therefore, it is more prudent to exclude the whole gene from downstream analysis.

Hence, *DESeq2* offers two possible responses to flagged outliers. By default, outliers in conditions with 6 or fewer replicates cause the whole gene to be flagged and removed from subsequent analysis, including *p*–value adjustment for multiple testing. For conditions that contain 7 or more replicates, *DESeq2* replaces the outlier counts with an imputed value, namely the trimmed mean over all samples, scaled by the size factor, and then re-estimates the dispersion, logarithmic fold changes and *p*–values for these genes. As the outlier is replaced with the value predicted by the null hypothesis of no differential expression, this is a more conservative choice than simply omitting the outlier. When there are many degrees of freedom, the second approach avoids discarding genes which might contain true differential expression.

Supplementary Figure S4 displays the outlier replacement procedure for a single gene in a 7 by 7 comparison of the Bottomly et al. [15] dataset. While the original fitted means are heavily influenced by a single sample with a large count, the corrected logarithmic fold changes provide a better fit to the majority of the samples.

### Regularized logarithm transformation

For certain analyses, it is useful to transform data to render them homoskedastic. As an example, consider the task of assessing sample similarities in an unsupervised manner using a clustering or ordination algorithm. For RNA-seq data, the problem of heteroskedasticity arises: if the data are given to such an algorithm on the original count scale, the result will be dominated by highly expressed, highly variable genes; if logarithm-transformed data are used, undue weight will be given to weakly expressed genes, which show exaggerated logarithmic fold changes, as discussed above. Therefore, we use the shrinkage approach of *DESeq2* to implement a “regularized logarithm” transformation (rlog), which behaves similarly to a log_2_ transformation for genes with high counts, while shrinking together the values for different samples for genes with low counts. It therefore avoids a commonly observed property of the standard logarithm transformation, the spreading apart of data for genes with low counts, where random noise is likely to dominate any biologically meaningful signal. When we consider the variance of each gene, computed across samples, these variances are stabilized–i.e., approximately the same, or homoskedastic-after the rlog transformation, while they would otherwise strongly depend on the mean counts. It thus facilitates multivariate visualization and ordinations such as clustering or principal component analysis that tend to work best when the variables have similar dynamic range.

**Figure 5.**
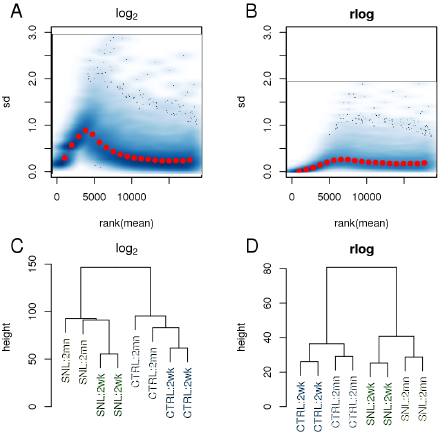
Variance stabilization and clustering after rlog transformation. Two transformations were applied to the counts of the Hammer et al. [25] dataset: the logarithm of normalized counts plus a pseudocount, i.e. *f(K_ij_)* = log_2_*(K_ij_/s_j_* + 1), and the rlog. The gene-wise standard deviation of transformed values is variable across the range of the mean of counts using the logarithm (A), while relatively stable using the rlog (B). A hierarchical clustering on Euclidean distances and complete linkage using the rlog (D) transformed data clusters the samples into the groups defined by treatment and time, while using the logarithm transformed counts (C) produces a more ambiguous result.

Note that while the rlog transformation builds upon on our LFC shrinkage approach, it is distinct from and not part of the statistical inference procedure for differential expression analysis described above, which employs the raw counts, not transformed data.

The rlog transformation is calculated by fitting for each gene a GLM with a baseline expression (i. e., intercept only) and, computing for each sample, shrunken logarithmic fold changes with respect to the baseline, using the same empirical Bayes procedure as before (Methods). Here, however, the sample covariate information (e. g. treatment or control) is not used, in order to treat all samples equally. The rlog transformation accounts for variation in sequencing depth across samples as it represents the logarithm of *q_ij_* after accounting for the size factors *s_ij_*. This is in contrast to the variance stabilizing transformation (VST) for overdispersed counts introduced in *DESeq* [4]: while the VST is also effective at stabilizing variance, it does not directly take into account differences in size factors; and in datasets with large variation in sequencing depth (dynamic range of size factors ≳ 4) we observed undesirable artifacts in the performance of the VST. A disadvantage of the rlog transformation with respect to the VST is, however, that the ordering of genes within a sample will change if neighboring genes undergo shrinkage of different strength. As with the VST, the value of rlog(*K_ij_*) for large counts is approximately equal to log_2_(*K_ij_/s_j_*). Both the rlog transformation and the VST are provided in the *DESeq2* package.

We demonstrate the use of the rlog transformation on the RNA-seq dataset of Hammer et al. [25], wherein RNA was sequenced from the dorsal root ganglion of rats which had undergone spinal nerve ligation and controls, at 2 weeks and at 2 months after the ligation. The count matrix for this dataset was downloaded from the ReCount online resource [26]. This dataset offers more subtle differences between conditions than the Bottomly et al. [15] dataset. Figure 5 provides diagnostic plots of the normalized counts under the ordinary logarithm with a pseudocount of 1 and the rlog transformation, showing that the rlog both stabilizes the variance through the range of the mean of counts and helps to find meaningful patterns in the data.

### Gene-level analysis

We here present *DESeq2* for the analysis of per-gene counts, i. e, the total number of reads that can be uniquely assigned to a gene. In contrast, several algorithms [27,28] work with probabilistic assignments of reads to transcripts, where multiple, overlapping transcripts can originate from each gene. It has been noted that the total read count approach can result in false detection of differential expression when in fact only transcript isoform lengths change, and even in a wrong sign of LFCs in extreme cases [27]. However, in our benchmark, discussed in the following section, we found that LFC sign disagreements between total read count and probabilistic assignment based methods were rare for genes that were differentially expressed according to either method (Supplementary Figure S5). Furthermore, if estimates for average transcript length are available for the conditions, these can be incorporated into the *DESeq2* framework as gene-and sample-specific normalization factors. In addition, the approach used in *DESeq2* can be extended to isoform-specific analysis, either through generalized linear modeling at the exon level with a gene-specific mean as in the *DEXSeq* package [29] or through counting evidence for alternative isoforms in splice graphs [30, 31]. In fact, the latest release version of *DEXSeq* now uses *DESeq2* as its inferential engine and so offers shrinkage estimation of dispersion and effect sizes for an exon-level analysis, too.

### Comparative benchmarks

To assess how well *DESeq2* performs for standard analyses in comparison to other current methods, we used a combination of simulations and real data. The Negative Binomial based approaches compared were *DESeq (old)* [4], *edgeR* [32], *edgeR* with the robust option [33], *DSS* [6] and *EBSeq* [34]. Other methods compared were the *voom* normalization method followed by linear modeling using the *limma* package [35] and the *SAMseq* permutation method of the *samr* package [23]. For the benchmarks using real data, the *Cuffdiff 2* [27] method of the Cufflinks suite was included.

For version numbers of the software used, see Supplementary Table S3. For all algorithms returning *p*–values, the *p*–values from genes with non-zero sum of read counts across samples were adjusted using the Benjamini-Hochberg procedure[20].

## Benchmarks through simulation

### Sensitivity and precision

We simulated datasets of 10,000 genes with Negative Binomial distributed counts. To simulate data with realistic moments, the mean and dispersions were drawn from the joint distribution of means and gene-wise dispersion estimates from the Pickrell et al. data, fitting only an intercept term. These datasets were of varying total sample size (*m* ∈ {6, 8,10, 20}), and the samples were split into two equal-sized groups. 80% of the simulated genes had no true differential expression, while for 20% of the genes, true fold changes of 2, 3, 4 were used to generate counts across the two groups, with the direction of fold change chosen randomly. The simulated DE genes were chosen uniformly at random among all the genes, throughout the range of mean counts. MA-plots of the true fold changes used in the simulation and the observed fold changes induced by the simulation for one of the simulation settings are shown in Supplementary Figure S6.

**Figure 6.**
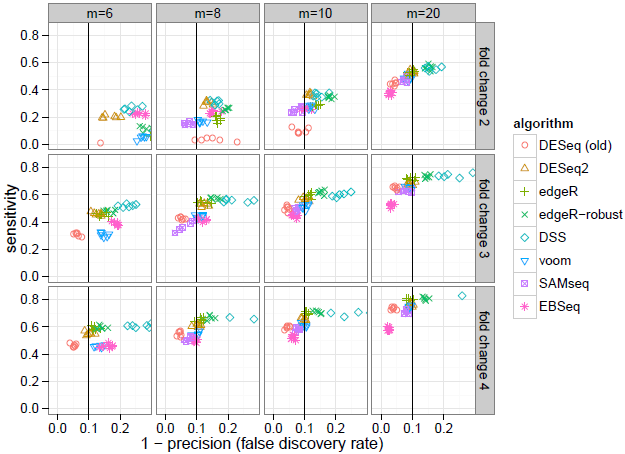
Sensitivity and precision of algorithms across combinations of sample size and effect size. *DESeq2* and *edgeR* often had the highest sensitivity of those algorithms which controlled the false discovery rate, i. e., those algorithms which fall on or to the left of the vertical black line. For a plot of sensitivity against false positive rate, rather than false discovery rate, see Supplementary Figure S8, and for the dependence of sensitivity on the mean of counts, see Supplementary Figure S9. Note that *EBSeq* filters low count genes (see main text for details).

Algorithms’ performance in the simulation benchmark was assessed by their sensitivity and precision. The sensitivity was calculated as the fraction of genes with adjusted *p*–value less than 0.1 among the genes with true differences between group means. The precision was calculated as the fraction of genes with true differences between group means among those with adjusted *p*–value less than 0.1. The sensitivity is plotted over 1 – precision, or the false discovery rate, in Figure 6. *DESeq2*, and also *edgeR*, often had the highest sensitivity of algorithms which controlled type-I error in the sense that the actual false discovery rate was at or below 0.1, the threshold for adjusted *p*–values used for calling differentially expressed genes. *DESeq2* revealed an increase in sensitivity over the other algorithms particularly for small fold change (fold change of 2 or 3), as was also found in benchmarks performed by Zhou et al. [33]. For larger sample sizes and larger fold changes the performance of the various algorithms was more consistent.

The overly conservative calling of the old *DESeq* tool can be observed, with reduced sensitivity compared to the other algorithms and an actual false discovery rate less than the nominal value of 0.1. We note that *EBSeq* version 1.4.0 by default removes low count genes – whose 75% quantile of normalized counts is less than 10 – before differential expression calling. The sensitivity of algorithms on the simulated data across a range of the mean of counts are more closely compared in Supplementary Figure S9.

### Outlier sensitivity

We used simulations to compare the sensitivity and specificity of *DESeq2*’s outlier handling approach to that of *edgeR*, which was recently added to the software and published while this manuscript was under review. *edgeR* now includes an optional method to handle outliers by iteratively refitting the generalized linear model after down-weighting potential outlier counts [33]. The simulations, summarized in Supplementary Figure S10, indicated that both approaches to outliers nearly recover the performance on an “outlier-free” dataset, though *edgeR-robust* had slightly higher actual than nominal false discovery rate, as seen in Supplementary Figure S11.

### Precision of fold change estimates

We benchmarked the *DESeq2* approach of using an empirical prior to achieve shrinkage of logarithmic fold change (LFC) estimates against two competing approaches: The *GFOLD* method, which allows for analysis of experiments without replication [19], can also handle experiments with replicates, and the *edgeR* package provides a pseudocount-based shrinkage termed *predictive logarithmic fold changes*. Results are summarized in Supplementary Figure S12–Figure S16. *DESeq2* had consistently low root mean squared error and mean absolute error across a range of sample sizes and models for the distribution of true logarithmic fold changes. *GFOLD* had similarly low error to *DESeq2* over all genes; however when focusing on the differentially expressed genes, it performed worse for larger sample sizes. *edgeR* with default settings had similarly low error to *DESeq2* when focusing only on the differentially expressed genes, but had higher error over all genes.

### Clustering

We compared the performance of the rlog transformation against other methods of transformation or distance calculation in the recovery of simulated clusters. The adjusted Rand Index [36] was used to compare a hierarchical clustering based on various distances with the true cluster membership. We tested the Euclidean distance on normalized counts, logarithm of normalized counts plus a pseudocount of 1, rlog transformed counts and VST counts. In addition we compared these Euclidean distances with the Poisson Distance implemented in the *PoiClaClu* package [37], and a distance implemented internally in the *plotMDS* function of *edgeR* (though not the default distance, which is similar to the logarithm of normalized counts). The results, shown in Supplementary Figure S17, revealed that when the size factors were equal for all samples, the Poisson Distance and the Euclidean distance of rlog-transformed or VST counts outperformed other methods. However, when the size factors were not equal across samples, the rlog approach generally outperformed the other methods. Finally, we note that the rlog transformation provides normalized data, which can be used for a variety of applications, of which distance calculation is one.

### Benchmark on RNA-seq data

While simulation is useful to verify how well an algorithm behaves with idealized, theoretical data, and hence allows verification that the algorithm performs as expected under its own assumptions, simulations cannot inform us how well the theory fits reality. With RNA-seq data, there is the complication of not knowing fully or directly the underlying truth; however, we can work around this limitation by using more indirect inference, explained below.

In the following benchmarks, we considered three performance metrics for differential expression calling: the false positive rate (or 1 minus the specificity), sensitivity and precision. We can obtain meaningful estimates of specificity from looking at datasets where we believe all genes fall under the null hypothesis of no differential expression [38]. Sensitivity and precision are more difficult to estimate, as they require independent knowledge of those genes that are differentially expressed. To circumvent this problem, we used experimental reproducibility on independent samples (though from the same dataset) as a proxy. We used a dataset with large numbers of replicates in both of two groups, where we expect that truly differentially expressed genes exist. We repeatedly split this dataset into an evaluation set and a larger verification set, and compared the calls from the evaluation set with the calls from the verification set, which were taken as “truth”. It is important to keep in mind that the calls from the verification set are only an approximation of the true differential state, and the approximation error has a systematic and a stochastic component. The stochastic error becomes small once the sample size of the verification set is large enough. For the systematic errors, our benchmark assumes that these affect all algorithms more or less equally and do not markedly change the ranking of the algorithms.

**Figure 7.**
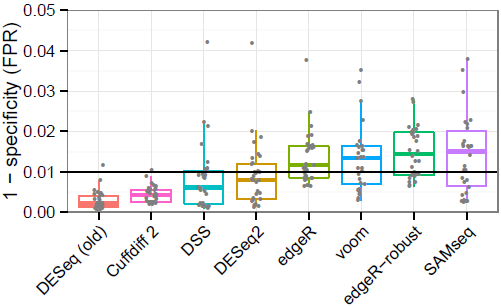
Benchmark of false positive calling. Shown are estimates of P(*p*– value < 0.01) under the null hypothesis. The number of *p*-values less than 0.01 divided by the total number of tests, from randomly selected comparisons of 5 vs 5 samples from the Pickrell et al. [16] dataset, with no known condition dividing the samples. Type-I error control requires that the tool does not substantially exceed the nominal value of 0.01 (black line). *EBSeq* results were not included in this plot as it returns posterior probabilities, which unlike *p*–values are not expected to be uniformly distributed under the null hypothesis.

### False positive rate

To evaluate the false positive rate of the algorithms, we considered mock comparisons from a dataset with many samples and no known condition dividing the samples into distinct groups. We used the RNA-seq data of Pickrell et al. [16] on lymphoblastoid cell lines derived from unrelated Nigerian individuals. We chose a set of 26 RNA-seq samples of the same read length (46 base pairs) and from male individuals. We randomly drew without replacement 10 samples from the set to perform a comparison of 5 against 5, and this process was repeated 30 times. We estimated the false positive rate associated with a critical value of 0.01 by dividing the number of *p*-values less than 0.01 by the total number of tests; genes with zero sum of read counts across samples were excluded. The results over the 30 replications, summarized in Figure 7, indicated that all algorithms generally controlled the number of false positives. *DESeq (old)* and *Cuffdiff 2* appeared overly conservative in this analysis, not using up their type-I error “budget”.

### Sensitivity

To obtain an impression of the sensitivity of the algorithms, we considered the Bottomly et al. [15] dataset, which contains 10 and 11 replicates of two different, genetically homogeneous mice strains. This allowed for a split of 3 vs 3 for the evaluation set and 7 vs 8 for the verification set, which were balanced across the 3 experimental batches. Random splits were replicated 30 times. Batch information was not provided to the *DESeq (old), *DESeq2*, DSS, edgeR*, and *voom* algorithms, which can accommodate complex experimental designs, in order to have comparable calls across all algorithms.

**Figure 8.**
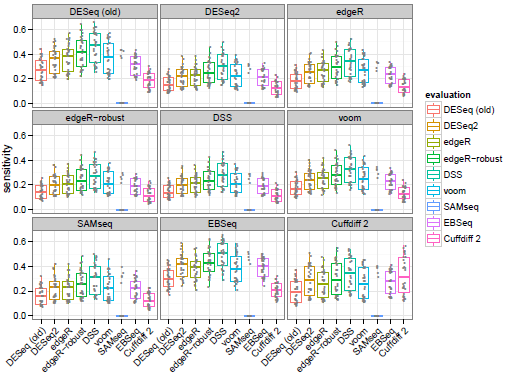
Sensitivity estimated from experimental reproducibility. Each algorithm’s sensitivity in the evaluation set (boxplots) is evaluated using the calls of each other algorithm in the verification set (panels with grey label).

We rotated though each algorithm to determine the calls of the verification set. Against a given algorithm’s verification set calls, we tested the evaluation set calls for every algorithm. We used this approach rather than a consensus-based method, as we did not want to favor or disfavor any particular algorithm or group of algorithms. Sensitivity was calculated as in the simulation benchmark, now with “true” differential expression defined by an adjusted *p*–value less than 0.1 in the larger verification set, as diagrammed in Supplementary Figure S18. Figure 8 displays the estimates of sensitivity for each algorithm pair.

The ranking of algorithms was generally consistent regardless of which algorithm was chosen to determine calls in the verification set. *DESeq2* had comparable sensitivity to *edgeR* and *voom* though less than *DSS*. The median sensitivity estimates were typically between 0.2 and 0.4 for all algorithms. That all algorithms had relatively low median sensitivity can be explained by the small sample size of the evaluation set and the fact that increasing the sample size in the verification set increases power. It is expected that the permutation-based *SAMseq* method rarely produced adjusted *p*–value less than 0.1 in the evaluation set, because the 3 vs 3 comparison does not enable enough permutations.

### Precision

Another important consideration from the perspective of an investigator is the precision, or fraction of true positives in the set of genes which pass the adjusted *p*–value threshold. This can also be reported as 1 – FDR, the false discovery rate. Again, “true” differential expression was defined by an adjusted *p*–value less than 0.1 in the larger verification set. The estimates of precision are displayed in Figure 9, where we can see that *DESeq2* often had the second highest median precision, behind *DESeq (old*). We can also see that algorithms which had higher median sensitivity, e.g., *DSS*, were generally associated here with lower median precision. The rankings differed significantly when *Cuffdiff 2* was used to determine the verification set calls. This is likely due to the additional steps *Cuffdiff 2* performed to deconvolve changes in isoform-level abundance from gene-level abundance, which apparently came at the cost of lowered precision when compared against its own verification set calls.

To further compare the sensitivity/precision results, we calculated the precision of algorithms along a grid of nominal adjusted *p*–values (Supplementary Figure S19). We then found the nominal adjusted *p*–value for each algorithm which resulted in a median actual precision of 0.9 (false discovery rate of 0.1). Having thus calibrated each algorithm to a target false discovery rate, we evaluated the sensitivity of calling, as shown in Supplementary Figure S20. As expected, here the algorithms performed more similarly to each other. This analysis revealed that, for a given target precision, *DESeq2* often was among the top algorithms by median sensitivity, though the variability across random replicates was larger than the differences between algorithms.

**Figure 9.**
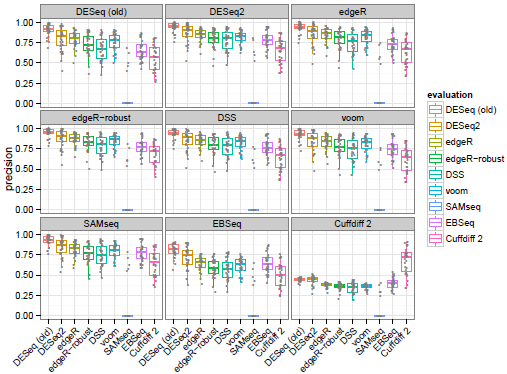
Precision estimated from experimental reproducibility. Each algorithm’s precision in the evaluation set (boxplots) is evaluated using the calls of each other algorithm in the verification set (panels with grey label).

The absolute number of calls for the evaluation and verification sets can be seen in Supplementary Figure S21 and Figure S22, which mostly matched the order seen in the sensitivity plot of Figure 8. Supplementary Figure S23 and Figure S24 provide heatmaps and clustering based on the Jaccard index of calls for one replicate of the evaluation and verification sets, indicating a large overlap of calls across the different algorithms.

In summary, the benchmarking tests showed that *DESeq2* effectively controlled type-I error, maintaining a median false positive rate just below the chosen critical value in a mock comparison of groups of samples randomly chosen from a larger pool. For both simulation and analysis of real data, *DESeq2* often achieved the highest sensitivity of those algorithms which controlled the false discovery rate.

## Conclusion

*DESeq2* offers a comprehensive and general solution for gene-level analysis of RNA-seq data. The use of shrinkage estimators substantially improves the stability and reproducibility of analysis results compared to maximum-likelihood based solutions. The use of empirical Bayes priors provides automatic control of the amount of shrinkage based on the amount of information for the estimated quantity available in the data. This allows *DESeq2* to offer consistent performance over a large range of data types and makes it applicable for small studies with few replicates as well as for large observational studies. *DESeq2*’s heuristics for outlier detection help to recognize genes for which the modeling assumptions are unsuitable and so avoids type-I errors caused by these. The embedding of these strategies in the framework of generalized linear models enables the treatment of both simple and complex designs.

A critical advance is the shrinkage estimator for fold changes for differential expression analysis, which offers a sound and statistically well-founded solution to the practically relevant problem of comparing fold change across the wide dynamic range of RNA-seq experiments. This is of value for many downstream analysis tasks, including the ranking of genes for follow-up studies and association of fold changes with other variables of interest. In addition, the rlog transformation, which implements shrinkage of fold changes on a per-sample basis, facilitates visualization of differences, for example in heatmaps, and enables the application of a wide range of techniques that require homoskedastic input data, including machine-learning or ordination techniques such as principal-component analysis and clustering.

*DESeq2* hence offers to practitioners a wide set of features with state-of-the-art inferential power. Its use cases are not limited to RNA-seq data or other transcriptomics assays; rather, many kinds of high-throughput count data can be used. Other areas for which *DESeq* or *DESeq2* have been used include ChIP-seq assays (e.g., [39]; see also the *DiffBind* package [40,41]), barcode-based assays (e.g., [42]), metagenomics data (e.g., [43]), ribosome profiling [44] and CRISPR/Cas-library assays [45]. Finally, the *DESeq2* package is well integrated in the Bioconductor infrastructure [10] and comes with extensive documentation, including a vignette that demonstrates a complete analysis step by step and discusses advanced use cases.

## Methods

A summary of the notation used in the following section is provided in Supplementary Table S1.

### Model and normalization

The read count *K_ij_* for gene *i* in sample *j* is described with a generalized linear model (GLM) of the Negative Binomial family with logarithmic link:

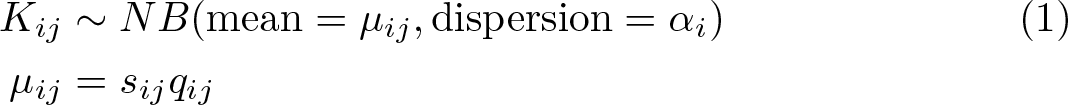

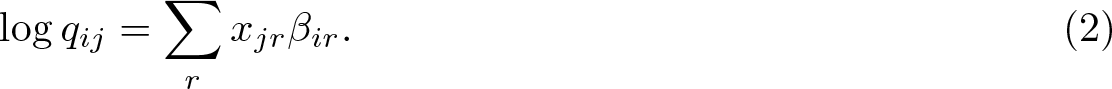

For notational simplicity, the equations here use the natural logarithm as the link function, though the *DESeq2* software reports estimated model coefficients and their estimated standard errors on the log_2_ scale.

By default, the normalization constants *s_ij_* are considered constant within a sample, *s_ij_* = *s_j_*, and are estimated with the median-of-ratios method previously described and used in *DESeq* [4] and *DEXSeq* [29]:

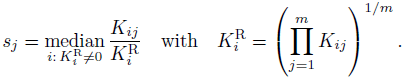

Alternatively, the user can supply normalization constants *s_ij_* calculated using other methods (e.g., using *cqn* [12] or *EDASeq* [13]), which may differ from gene to gene.

### Expanded design matrices

For consistency with our software’s documentation, in the following text we will use the terminology of the *R* statistical language. In linear modeling, a categorical variable or *factor* can take on two or more values or *levels*. In standard design matrices, one of the values is chosen as a reference value or *base level* and absorbed into the intercept. In standard GLMs, the choice of base level does not inuence the values of contrasts (LFCs). This, however, is no longer the case in our approach using ridge-regression-like shrinkage on the coeffcients (described below), when factors with more than two levels are present in the design matrix, because the base level will not undergo shrinkage while the other levels do.

To recover the desirable symmetry between all levels, *DESeq2 uses expanded design matrices* which include an indicator variable for each level of each factor, in addition to an intercept column (i. e., none of the levels is absorbed into the intercept). While such a design matrix no longer has full rank, a unique solution exists because the zero-centered prior distribution (see below) provides regularization. For dispersion estimation and for estimating the width of the LFC prior, standard design matrices are used.

### Contrasts

Contrasts between levels and standard errors of such contrasts can be calculated as they would in the standard design matrix case, i. e., using:

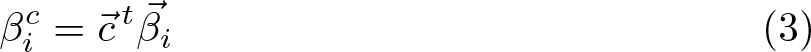

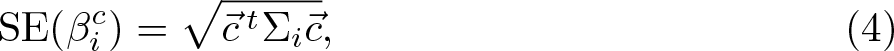

where 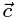 represents a numeric contrast, e.g., 1 and −1 specifying the numerator and denominator of a simple two level contrast, and 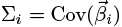, defined below.

### Estimation of Dispersions

We assume the dispersion parameter *α_1_* follows a log-Normal prior distribution that is centered around a trend that depends on the gene’s mean normalized read count:

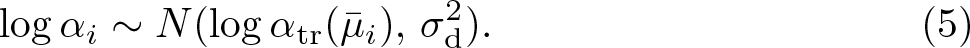

Here, *α_tr_* is a function of the gene’s mean normalized count, 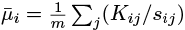. It describes the mean-dependent expectation of the prior. *σ*_d_ is the width of the prior, a hyperparameter describing how much the individual genes' true dispersions scatter around the trend. For the trend function, we use the same parametrization as we used for *DEXSeq* [29], namely,

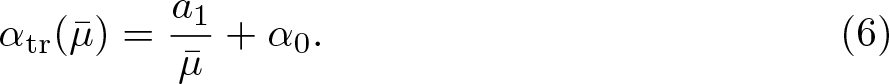

We get final dispersion estimates from this model in three steps, which implement a computationally fast approximation to a full empirical Bayes treatment. We first use the count data for each gene separately to get preliminary gene-wise dispersion estimates 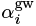 by maximum likelihood estimation. Then, we fit the dispersion trend *α*_tr_. Finally, we combine the likelihood with the trended prior to get maximum *a posteriori* (MAP) values as final dispersion estimates. Details for the three steps follow.

### Gene-wise dispersion estimates

To get a gene-wise dispersion estimate for a gene *i*, we start by fitting a Negative Binomial GLM without logarithmic fold change prior for the design matrix *X* to the gene’s count data. This GLM uses a rough method-of-moments estimate of dispersion, based on the within-group variances and means. The initial GLM is necessary to obtain an initial set of fitted values, 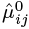. We then maximize the Cox-Reid adjusted likelihood of the dispersion, conditioned on the fitted values 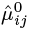 from the initial fit, to obtain the gene-wise estimate 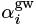, i.e.,

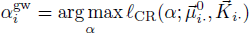

with

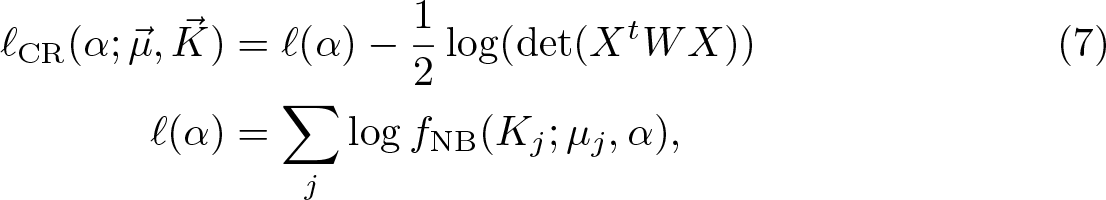

where *f*_NB_(*k*; *μ α*) is the probability mass function of the Negative Binomial distribution with mean *μ* and dispersion *α*, and the second term provides the Cox-Reid bias adjustment [46]. This adjustment, first used in the context of dispersion estimation for SAGE data [47] and then for HTS data [3] in *edgeR*, corrects for the negative bias of dispersion estimates from using the maximum likelihood estimates (MLE) for the fitted values 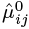 (analogous to Bessel’s correction in the usual sample variance formula; for details, see [48, Section 10.6]). It is formed from the Fisher information for the fitted values, which is here calculated as det(*X^t^WX*), where *W* is the diagonal weight matrix from the standard iteratively re-weighted least squares (IRLS) algorithm. As the GLM’s link function is *g(μ)* = log(*μ*) and its variance function is *V*(*μ*; *α*)= *μ* + *α μ^2^*, the elements of the diagonal matrix *W_i_* are given by:

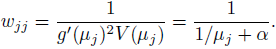

The optimization in Equation (7) is performed on the scale of log *α* using a backtracking line search with proposals accepted which satisfy Armijo conditions [49].

### Dispersion trend

A parametric curve of the form (6) is fit by regressing the gene-wise dispersion estimates 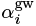 onto the means of the normalized counts, 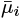. The sampling distribution of the gene-wise dispersion estimate around the true value *α_i_* can be highly skewed, and therefore we do not use ordinary least square regression but rather Gamma-family GLM regression. Furthermore, dispersion outliers could skew the fit and hence a scheme to exclude such outliers is used.

The hyperparameters *a_1_* and *α_0_* of (6) are obtained by iteratively fitting a Gamma-family GLM. At each iteration, genes with ratio of dispersion to fitted value outside the range [10^−4^,15] are left out until the sum of squared logarithmic fold changes of the new coefficients over the old coefficients is less than 10^−6^ (same approach as in *DEXSeq* [29]).

The parametrization (6) is based on reports by us and others of decreasing dependence of dispersion on the mean in many datasets [4,5,50,3,6]. Some caution is warranted to disentangle true underlying dependence from effects of estimation bias that can create a perceived mean-dependence of the dispersion. Consider a Negative Binomial distributed random variable with expectation μ and dispersion *α*. Its variance *v* = *μ*+ *αμ*^2^ has two components, *v* = *v_P_* +*v_D_*, the Poisson component *v_P_* = *μ* independent of *α* and the overdispersion component *v_D_* = *αμ*^2^. When *μ* is small,(*μ* ≲ 1/*α*) (vertical lines in Supplementary Figure S1),the Poisson component dominates, in the sense that *v_P_/v_D_* = 1/(*αμ*)≳1, and the observed data provide little information on the value of *α*. Therefore the sampling variance of an estimator for *α* will be large when *μ* ≲ 1/*α*, which leads to the appearance of bias. For simplicity, we have stated the above argument without regard to the influence of the size factors, *s_j_*, on the value of *μ*. This is permissible because, by construction, the geometric mean of our size factors is close to 1, and hence, the mean across samples of the unnormalized read counts, 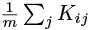, and the mean of the normalized read counts, 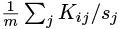, will be roughly the same.

This phenomenon may give rise to an apparent dependence of *α* on *μ*. It is possible that the shape of the dispersion-mean fit for the Bottomly data (Figure 1A) can be explained in that manner: the asymptotic dispersion is *α_0_* ≈ 0.01, and the non-zero slope of the mean-dispersion plot is limited to the range of mean counts up to around 100, the reciprocal of *α_0_*. However, overestimation of *α* in that low-count range has little effect on inference, as in that range the variance *v* is anyway dominated by the *α*-independent Poisson component *v_P_*. The situation is different for the Pickrell data: here, a dependence of dispersion on mean was observed for counts clearly above the reciprocal of the asymptotic dispersion *α_0_* (Figure 1B), and hence is not due merely to estimation bias. Simulations (shown in Supplementary Figure S25) confirmed that the observed joint distribution of estimated dispersions and means is not compatible with a single, constant dispersion. Therefore, the parametrization (6) is a flexible and mildly conservative modeling choice: it is able to pick up dispersion-mean dependence if it is present, while it can lead to a minor loss of power in the low count range due to a tendency to overestimate dispersion there.

### Dispersion prior

As also observed by Wu et al. [6], a log-Normal prior fits the observed dispersion distribution for typical RNA-seq datasets. We solve the computational difficulty of working with a non-conjugate prior using the following argument: the logarithmic residuals from the trend fit, 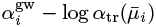, arise from two contributions, namely the scatter of the true logarithmic dispersions around the trend, given by the prior with variance 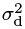, and the sampling distribution of the logarithm of the dispersion estimator, with variance 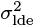. The sampling distribution of a dispersion estimator is approximately a scaled *χ*^2^ distribution with *m* – *p* degrees of freedom, with *m* the number of samples and *p* the number of coefficients. The variance of the logarithm of 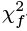–distributed random variable is given [51] by the trigamma function *ψ*_1_,

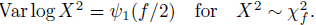

Therefore, 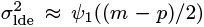, i.e., the sampling variance of the logarithm of a variance or dispersion estimator is approximately constant across genes and depends only on the degrees of freedom of the model.

Supplementary Table S2 compares this approximation for the variance of logarithmic dispersion estimates with the variance of logarithmic Cox-Reid adjusted dispersion estimates for simulated Negative Binomial data, over a combination of different sample sizes, number of parameters and dispersion values used to create the simulated data. The approximation is close to the sample variance for various typical values of *m*, *p* and *α*.

Therefore, the prior variance 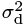 is obtained by subtracting the expected sampling variance from an estimate of the variance of the logarithmic residuals, 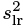:

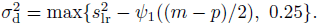

The prior variance 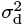 is thresholded at a minimal value of 0.25 so that the dispersion estimates are not shrunk entirely to 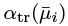 in the case that the variance of the logarithmic residuals is less than the expected sampling variance.

To avoid inflation of 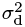 due to dispersion outliers (i. e., genes not well captured by this prior; see below), we use a robust estimator for the standard deviation *s_Ir_* of the logarithmic residuals,

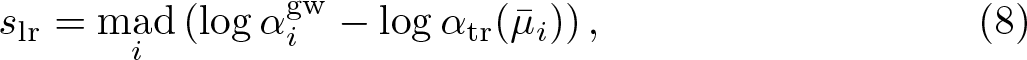

where mad stands for the median absolute deviation, divided as usual by the scaling factor Φ^−1^(3/4).

### Three or less residuals degrees of freedom

When there are 3 or less residual degrees of freedom (number of samples minus number of parameters to estimate), the estimation of the prior variance 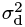 using the observed variance of logarithmic residuals 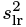 tends to underestimate 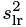. In this case, we instead estimate the prior variance through simulation. We match the distribution of logarithmic residuals to a density of simulated logarithmic residuals. These are the logarithm of 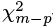–distributed random variables added to 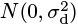 random variables to account for the spread due to the prior. The simulated distribution is shifted by – log(*m* – *p*) to account for the scaling of the *χ*^2^ distribution. We repeat the simulation over a grid of values for 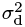, and select the value which minimizes the Kullback-Leibler divergence from the observed density of logarithmic residuals to the simulated density.

### Final Dispersion Estimate

We form a logarithmic posterior for the dispersion from the Cox-Reid adjusted logarithmic likelihood (7) and the logarithmic prior (5) and use its maximum (i. e., the maximum *a posteriori*, MAP, value) as the final estimate of the dispersion,

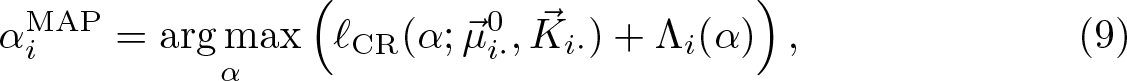

where

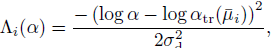

is, up to an additive constant, the logarithm of the density of prior (5). Again, a backtracking line search is used to perform the optimization.

### Dispersion outliers

For some genes, the gene-wise estimate 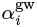 can be so far above the prior expectation 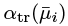 that it would be unreasonable to assume that the prior is suitable for the gene. If the dispersion estimate for such genes were down-moderated toward the fitted trend, this might lead to false positives. Therefore, we use the heuristic of considering a gene as a “dispersion outlier”, if the residual from the trend fit is more than two standard deviations of logarithmic residuals, s_lr_ (see Equation (8)), above the fit, i.e., if

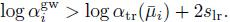

For such genes, the gene-wise estimate 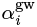 is not shrunk toward the trended prior mean. Instead of the MAP value 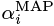, we use the gene-wise estimate 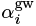 as a final dispersion value in the subsequent steps. In addition, the iterative fitting procedure for the parametric dispersion trend described above avoids that such dispersion outliers influence the prior mean.

## Shrinkage estimation of logarithmic fold changes

To incorporate empirical Bayes shrinkage of logarithmic fold changes, we postulate a zero-centered Normal prior for the coefficients *β_ir_* of model (2) that represent logarithmic fold changes (i.e., typically, all coefficients except for the intercept *β_ir_*):

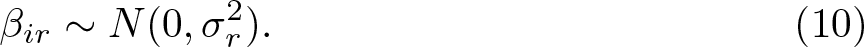

As was observed with differential expression analysis using microarrays, genes with low intensity values tend to suffer from a small signal-to-noise ratio. Alternative estimators can be found which are more stable than the standard calculation of fold change as the ratio of average observed values for each condition [52,53,54]. *DESeq2*’s approach can be seen as an extension of these approaches for stable estimation of gene expression fold changes to count data.

### Empirical prior estimate

To obtain values for the empirical prior widths *σ_r_* for the model coefficients, we again approximate a full empirical Bayes approach, as with the estimation of dispersion prior, though here we do not subtract the expected sampling variance from the observed variance of maximum likelihood estimates. The estimate of the logarithmic fold change prior width is calculated as follows. We use the standard iteratively reweighted least squares (IRLS) algorithm [11] for each gene’s model (1,2) to get maximum likelihood estimates for the coefficients 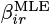. We then fit, for each column r of the design matrix (except for the intercept) a zero-centered Normal to the empirical distribution of MLE fold change estimates 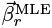.

To make the fit robust against outliers with very high absolute LFC values, we use quantile matching: the width *σ_r_* is chosen such that the (1 – *p*) empirical quantile of the absolute value of the observed LFCs, 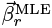, matches the (1 – *p*/2) theoretical quantile of the prior, 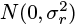, where *p* is set by default to 0.05. If we write the theoretical upper quantile of a Normal distribution as Q_N_ (1 – *p*) and the empirical upper quantile of the MLE LFCs as Q_| βr|_(1 – *p*), then the prior width is calculated as:

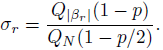

To ensure that the prior width *σ_r_* will be independent of the choice of base level, the estimates from the quantile matching procedure are averaged for each factor over all possible contrasts of factor levels. When determining the empirical upper quantile, extreme LFC values (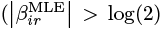, or 10 on the base 2 scale) are excluded.

### Final estimate of logarithmic fold changes

The logarithmic posterior for the vector, 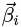, of model coefficients *β*_*ir*_ for gene *i* is the sum of the logarithmic likelihood of the GLM (2) and the logarithm of the prior density (10), and its maximum yields the final MAP coefficient estimates:

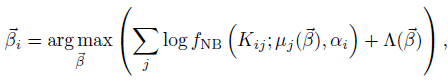

where

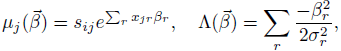

and α_*i*_. is the final dispersion estimate for gene *i*, i.e., 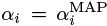, except for dispersion outliers, where 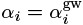.

The term ∧(*β*), i. e., the logarithm of the density of the Normal prior (up to an additive constant), can be read as a ridge penalty term, and therefore, we perform the optimization using the *iteratively reweighted ridge regression algorithm* [55], also known as *weighted updates* [56]. Specifically, the updates for a given gene are of the form

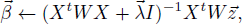

with 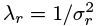 and

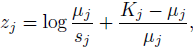

where the current fitted values 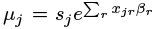,are computed from the current estimates 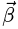 in each iteration.

### Fisher information

The effect of the zero-centered Normal prior can be understood as to shrink the MAP LFC estimates based on the amount of information the experiment provides for this coefficient, and we briefly elaborate on this here. Specifically, for a given gene *i*, the shrinkage for an LFC *β_ir_* depends on the *observed Fisher information*, given by

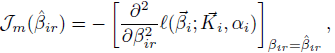

where 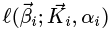; is the logarithm of the likelihood, and partial derivatives are taken with respect to LFC β_*ir*_. For a Negative Binomial GLM, the observed Fisher information, or peakedness of the logarithm of the profile likelihood, is influenced by a number of factors including the degrees of freedom, the estimated mean counts *μ_ij_*, and the gene’s dispersion estimate *α_i_*. The prior exerts its influence on the MAP estimate when the density of the likelihood and the prior are multiplied to calculate the posterior. Genes with low estimated mean values *μ_ij_* or high dispersion estimates a. have flatter profile likelihoods, as do datasets with few residual degrees of freedom, and therefore in these cases the zero-centered prior pulls the MAP estimate from a high-uncertainty MLE closer toward zero.

## Wald test

The Wald test compares the beta estimate *β_ir_* divided by its estimated standard error SE(*β_ir_*) to a standard Normal distribution. The estimated standard errors are the square root of the diagonal elements of the estimated covariance matrix, ∑_*i*_, for the coefficients, i. e., 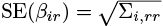. Contrasts of coefficients are tested similarly by forming a Wald statistics using (3) and (4). We use the following formula for the coefficient covariance matrix for a generalized linear model with Normal prior on coefficients [55,57]:

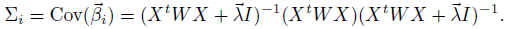

The tail integrals of the standard Normal distribution are multiplied by 2 to achieve a two-tailed test. The Wald test *p*–values from the subset of genes which pass the independent filtering step are adjusted for multiple testing using the procedure of Benjamini and Hochberg [[20].

## Independent filtering

Independent filtering does not compromise type-I error control as long as the distribution of the test statistic is marginally independent of the filter statistic *under the null hypothesis* [21], and we argue in the following that this is the case in our application. The filter statistic in *DESeq2* is the mean of normalized counts for a gene, while the test statistic is *p*, the *p*–value from the Wald test. We first consider the case where the size factors are equal and where the gene-wise dispersion estimates are used for each gene, i.e. without dispersion shrinkage. The distribution family for the Negative Binomial is parameterized by θ = (μ, α). Aside from discreteness of *p* due to low counts, for a given μ, the distribution of *p* is Uniform(0,1) under the null hypothesis, so *p* is an ancillary statistic. The sample mean of counts for gene *i*, 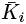, is boundedly complete sufficient for μ. Then from Basu’s theorem, we have that 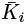. and *p* are independent.

While for very low counts, one can observe discreteness and non-uniformity of *p* under the null, *DESeq2* does not use the distribution of the *p* in its estimation procedure – for example, *DESeq2* does not estimate the proportion of null genes using the distribution of *p* – so this kind of dependence of *p* on *μ* does not lead to increased type-I error.

If the size factors are not equal across samples, but not correlated with condition, conditioning on the mean of *normalized* counts should also provide uniformly distributed *p* as with conditioning on the mean of counts, 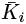. We may consider a pathological case where the size factors are perfectly confounded with condition, in which case, even under the null hypothesis, genes with low mean count would have non-uniform distribution of *p*, as one condition could have positive counts and the other condition often zero counts. This could lead to non-uniformity of *p* under the null hypothesis, however such a pathological case would pose problems for many statistical tests of differences in mean.

We used simulation to demonstrate that the independence of the null distribution of the test statistic from the filter statistic still holds in the case of dispersion shrinkage. Supplementary Figure S26 displays marginal null distributions of *p* across the range of mean normalized counts. Despite spikes in the distribution for the genes with the lowest mean counts due to discreteness of the data, these densities were nearly uniform across the range of average expression strength.

## Composite null hypotheses

*DESeq2* offers tests for composite null hypotheses of the form 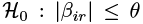 in order to find genes whose LFC significantly exceeds a threshold θ > 0. The composite null hypothesis is replaced by two simple null hypotheses: 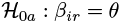 and 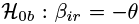. Two-tailed p-values are generated by integrating a Normal distribution centered on *θ* with standard deviation SE(β_*ir*_) from |β_*ir*_| toward ∞. The value of the integral is then multiplied by 2 and thresholded at 1. This procedure controls type-I error even when *β_ir_* = ± θ, and is equivalent to the standard *DESeq2 p*–value when θ = 0.

Conversely, when searching for genes whose absolute LFC is significantly below a threshold, i.e., when testing the null hypothesis 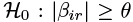, the *p*–value is constructed as the maximum of two one-sided tests of the simple null hypotheses: 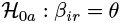 and 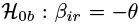. The one-sided *p*–values are generated by integrating a Normal distribution centered on θ with standard deviation SE(*β_ir_*) from *β_ir_* toward −∞, and integrating a Normal distribution centered on −θ with standard deviation SE(*β_ir_*) from *β_ir_* toward ∞.

Note that while a zero-centered prior on LFCs is consistent with testing the null hypothesis of small LFCs, it should not be used when testing the null hypothesis of large LFCs, because the prior would then favor the alternative hypothesis. *DESeq2* requires that no prior has been used when testing the null hypothesis of large LFCs, so that the data alone must provide evidence against the null.

## Interactions

Two exceptions to the default *DESeq2* LFC estimation steps are used in the case of experimental designs with interaction terms. First, when any interaction terms are included in the design, the LFC prior width for main effect terms is not estimated from the data, but set to a wide value (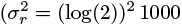, or 1000 on the base 2 scale). This ensures that shrinkage of main effect terms will not result in false positive calls of significance for interactions. Second, when interaction terms are included and all factors have two levels, then standard design matrices are used rather than expanded model matrices, such that only a single term is used to test the null hypothesis that a combination of two effects is merely additive in the logarithmic scale.

## Regularized logarithm

The rlog transformation is calculated as follows. The experimental design matrix *X* is substituted with a design matrix with an indicator variable for every sample in addition to an intercept column. A model as described in Equation (1),Equation (2) is fit with a zero-centered Normal prior on the non-intercept terms and using the fitted dispersion values 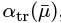, which capture the overall variance-mean dependence of the dataset. The true experimental design matrix *X* is then only used in estimating the variance-mean trend over all genes. For the purpose of unsupervised analyses, for instance sample quality assessment, it is desirable that the experimental design have no influence on the transformation, and hence *DESeq2* by default ignores the design matrix and re-estimates the dispersions treating all samples as replicates, i. e., uses *blind* dispersion estimation. The rlog transformed values are the fitted values,

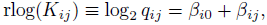

where *β_ij_* is the shrunken LFC on the base 2 scale for the *j*–th sample. The variance of the prior is set using a similar approach as taken with differential expression, by matching a zero-centered Normal distribution to observed LFCs. First a matrix of logarithmic fold changes is calculated by taking the logarithm (base 2) of the normalized counts plus a pseudocount of ½ for each sample divided by the mean of normalized counts plus a pseudocount of ½. The pseudocount of ½ allows for calculation of the logarithmic ratio for all genes, and has little effect on the estimate of the variance of the prior or the final rlog transformation. This matrix of LFCs then represents the common-scale logarithmic ratio of each sample to the fitted value using only an intercept. The prior variance is found by matching the 95% quantile of a zero-centered Normal distribution to the 90% quantile of the absolute values in the logarithmic fold change matrix.

## Cook’s distance for outlier detection

The maximum likelihood estimate of 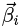 is used for calculating Cook’s distance. Considering a gene *i* and sample *j*, Cook’s distance for generalized linear models is given by [58]:

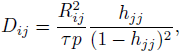

where *R_ij_* is the Pearson residual of sample *j*, *τ* is an overdispersion parameter (in the Negative Binomial GLM, τ is set to 1), *p* is the number of parameters including the intercept, and *h_jj_*, is the *j*–th diagonal element of the hat matrix ***H***:

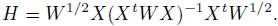

Pearson residuals *R_ij_* are calculated as

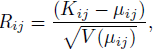

where μ_*ij*_ is estimated by the Negative Binomial GLM without the logarithmic fold change prior, and using the variance function *V*(μ) = μ+±μ^2^. A method of moments estimate 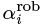, using a robust estimator of variance 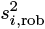 to provide robustness against outliers, is used here:

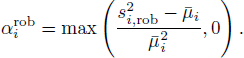

## R/Bioconductor package

*DESeq2* is implemented as a package for the R statistical environment and available as part of the Bioconductor project [10] at http://www.bioconductor.org/packages/release/bioc/html/DESeq2.html. The count matrix and metadata including the gene model and sample information are stored in an S4 class derived from the *SummarizedExperiment* class of the *GenomicRanges* package [59]. *SummarizedExperiment* objects containing count matrices can be easily generated using the *summarizeOverlaps* function of the *GenomicAlignments* package [60]. This workflow automatically stores the gene model as metadata and additionally other information such as the genome and gene annotation versions. Other methods to obtain count matrices include the *htseq-count* script [61] and the Bioconductor packages *easyRNASeq* [62] and *featureCount* [63].

The *DESeq2* package comes with a detailed vignette working through a number of example differential expression analyses on real datasets, and the use of the rlog transformation for quality assessment and visualization. A single function, called *DESeq*, is used to run the default analysis, while lower-level functions are also available for advanced users.

## Read Alignment for the Bottomly et al. and Pickrell et al. Datasets

Reads were aligned using the TopHat2 aligner [64], and assigned to genes using the *summarizeOverlaps* function of the *GenomicRanges* package [59]. The Sequence Read Archive fastq files of the Pickrell et al. [16] dataset (accession number [SRP001540]) were aligned to the Homo Sapiens reference sequence GRCh37 downloaded March 2013 from Illumina iGenomes. Reads were counted in the genes defined by the Ensembl GTF file, release 70, contained in the Illumina iGenome. The Sequence Read Archive fastq files of the Bottomly et al. [15]
dataset (accession number [SRP004777]) were aligned to the Mus Musculus reference sequence NCBIM37 downloaded March 2013 from Illumina iGenomes. Reads were counted in the genes defined by the Ensembl GTF file, release 66, contained in the Illumina iGenome.

## Reproducible code

Sweave vignettes for reproducing all figures and tables in this paper, including data objects for the experiments mentioned, and code for aligning reads and for benchmarking, can be found in a package *DESeq2paper*, available at http://www-huber.embl.de/DESeq2paper.

## List of Abbreviations

FDR: false discovery rate
GLM: generalized linear model
HTS: high-throughput sequencing
LFC: logarithmic fold change
MAP: maximum *a posteriori*
MLE: maximum likelihood estimate
SE: standard error
VST: variance-stabilizing transformation

## Author’s contributions

All authors developed the method and wrote the manuscript. MIL implemented the method and performed the analyses.

## Acknowledgements

The authors thank all users of DESeq and *DESeq2* who provided valuable feedback. We thank Judith Zaugg for helpful comments on the manuscript. ML acknowledges funding via a stipend from the International Max Planck Research School for Computational Biology and Scientific Computing and a grant from the National Institutes of Health (5T32CA009337-33). WH and SA acknowledge funding from the European Union’s 7th Framework Programme (Health) via Project *Radiant*. We thank an anonymous reviewer for raising the question of estimation biases in the dispersion-mean trend fitting.

## Supplement

### Supplementary Methods

#### Benchmarking Code

The code used to run the count-based algorithms is contained in the file inst/ script/runScripts.R in the *DESeq2paper* package (available at http://www.huber.embl.de/DESeq2paper). The code for the simulations is referenced from the simulations vignette in this package. The code which ran the algorithms over the real datasets is contained in the files inst/script/pickrell/diffExpr.R (the specificity analysis run on the Pickrell et al. [16] dataset) and inst/script/ bottomly/diffExpr.R (for the sensitivity and precision analysis run on the Bottomly et al. [15] dataset). The *Cuffdiff 2* commands are contained in the inst/ script/pickrell/ and inst/script/bottomly/ directories.

### Supplementary Tables

**Supplementary Table S1.**
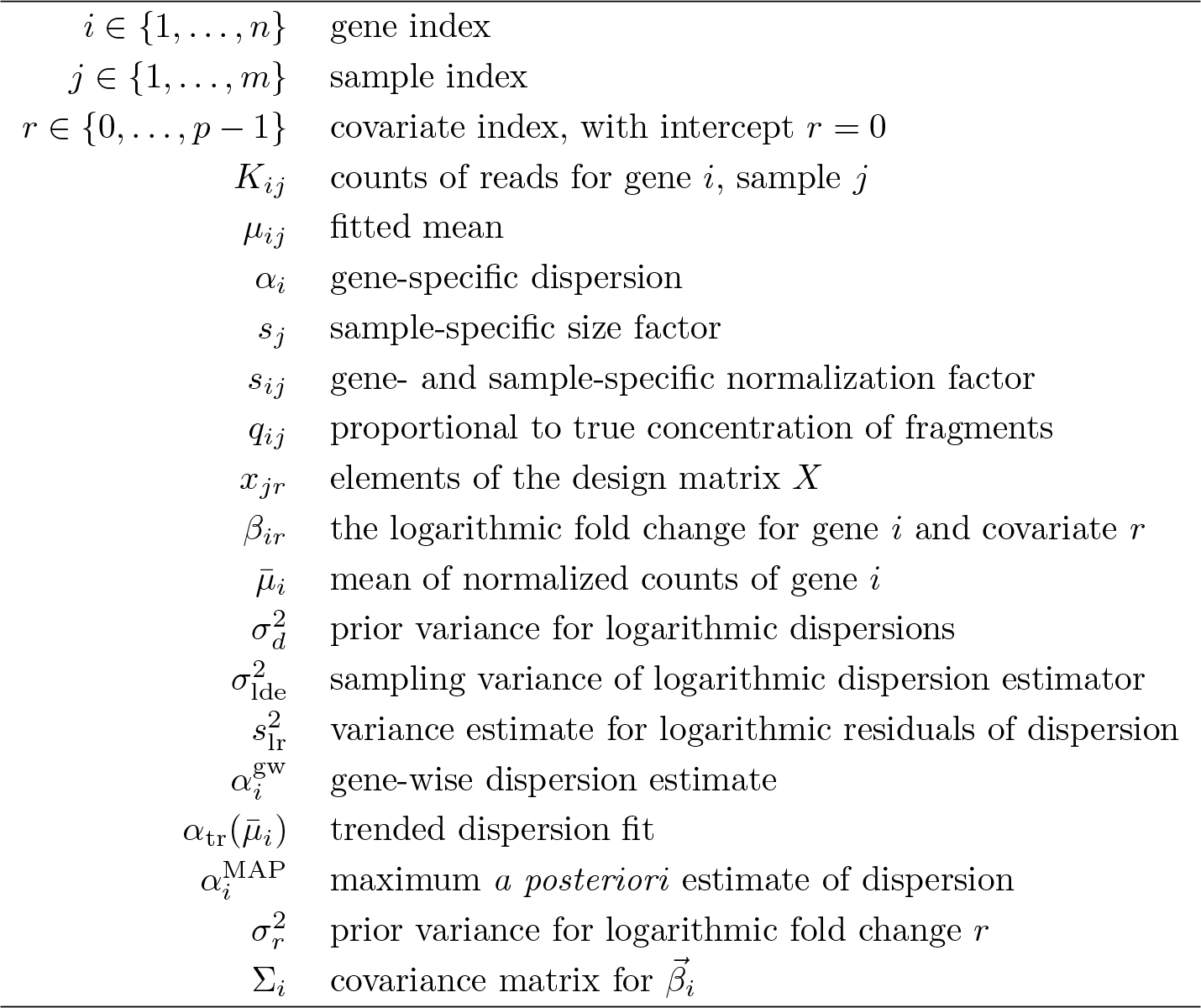
Notation

**Supplementary Table S2.** Theoretical and sample variance of logarithmic dispersion estimates for various combinations of sample size m, number of parameters p and true dispersion α. The estimates are the *DESeq2* gene-wise estimates from 4000 simulated genes with Negative Binomial counts with a mean of 1024. The sample variance of the logarithmic dispersion estimates is generally close to the approximation of theoretical variance.

| m | p | $\alpha$ | theor. var. | sample var. |
| --- | --- | --- | --- | --- |
| 6 | 2 | 0.05 | 0.645 | 0.670 |
| 6 | 2 | 0.20 | 0.645 | 0.642 |
| 8 | 2 | 0.05 | 0.395 | 0.409 |
| 8 | 2 | 0.20 | 0.395 | 0.396 |
| 8 | 3 | 0.05 | 0.490 | 0.530 |
| 8 | 3 | 0.20 | 0.490 | 0.462 |
| 16 | 2 | 0.05 | 0.154 | 0.160 |
| 16 | 2 | 0.20 | 0.154 | 0.138 |
| 16 | 3 | 0.05 | 0.166 | 0.169 |
| 16 | 3 | 0.20 | 0.166 | 0.156 |

**Supplementary Table S3.** Versions of software used in manuscript.

| function/package | version | additional information |
| --- | --- | --- |
| *DESeq (old)* | 1.16.0 | using the GLM test |
| *DESeq2* | 1.4.0 |  |
| *edgeR* | 3.6.0 | using GLM and trended dispersion estimation |
| *DSS* | 2.2.0 |  |
| *voom: limma* | 3.20.1 |  |
| *SAMseq: samr* | 2.0 | using `samr.pvalues.from.perms` for $p$-values |
| *EBSeq* | 1.4.0 | (1 – PPDE) used for FDR cutoff, following user guide |
| *Cuffdiff 2* | 2.1.1 |  |
| *GFOLD* | 1.1.2 |  |
| *PoiClaClu* | 1.0.2 |  |

### Supplementary Figures

**Supplementary Figure S1:**
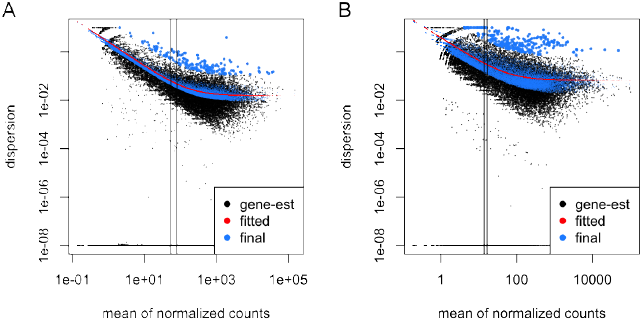
Shrinkage estimation of dispersion over all genes. Plot of dispersion estimates over the average expression strength (A) for the Bottomly et al. [15] dataset with 6 samples across 2 conditions and (B) for the Pickrell et al. [16] dataset with 5 samples fitting only an intercept term. This plot shows the same data as Figure 1, but with dispersions drawn for all genes instead of only a subset. The points at the bottom of the plot typically arise from genes for which the observed variance is below the variance expected under a Poisson model. In such a case, the maximum-likelihood estimate will be essential zero, and appears here with the surrogate value 10^−8^. Vertical lines indicate the reciprocal of the asymptotic dispersion a0, on the scale of raw counts for the samples with the smallest and largest size factor. The lines hence mark the count range where Poisson noise and overdispersion contribute about equally to the observed variance. For very low count values (left of the lines), dispersion estimates become unreliable, causing possible overestimation (Methods).

**Supplementary Figure S2:**
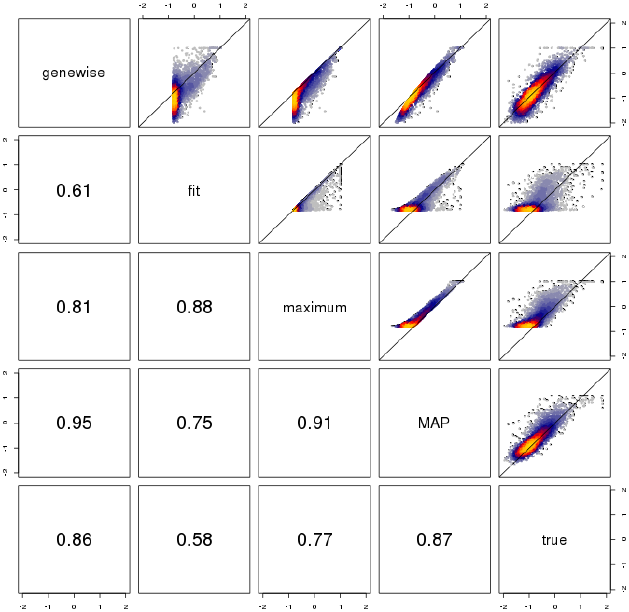
Scatterplot of various estimates of dispersion using *DESeq2*, against the true dispersion in the logarithmic scale (base 10) from simulated counts. The blue, red, and yellow colors indicate regions of increasing density of points. Counts for 4000 genes and for 10 samples in two groups were simulated with no true difference in means. The Negative Binomial counts had mean and dispersion drawn from the joint distribution of the mean and gene-wise dispersion estimates from the Pickrell et al. dataset. The estimates shown are *genewise*, the CR-adjusted maximum likelihood estimate; *fit* the value from the fitted curve; *maximum*, the maximum of the two previous values (the estimate used in the older version of *DESeq*); and *MAP*, the maximum *a posteriori* estimate used in *DESeq2*. The correlations shown in the bottom panels do not include the very low gene-wise estimates of dispersion which can result in potential false positives. The *MAP*, shrunken estimates used in *DESeq2* were closer to the diagonal, while the *maximum* estimate was typically above the true value of dispersion, which can lead to overly-conservative inference of differential expression.

**Supplementary Figure S3:**
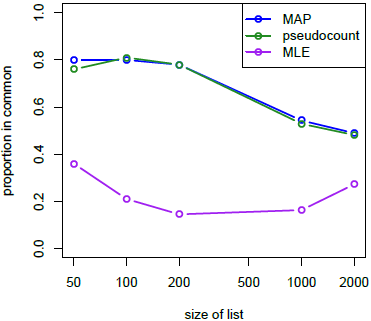
“Concordance at the top” plot. *DESeq2* is run on equally split halves of the data of Bottomly et al. [15] and the proportion of genes in common after ranking by absolute logarithmic fold changes is compared [65]. On the y-axis is the number of genes in common between the splits divided by the size of the top-ranked list. The MAP estimate of logarithmic fold change and the MLE after adding a pseudocount of 1 to all samples provide nearly the same concordance for various cutoffs, while ranking by the MLE on raw counts has generally low concordance. For further demonstrations of the advantage of MAP over pseudocount, see section *Benchmarks*.

**Supplementary Figure S4:**
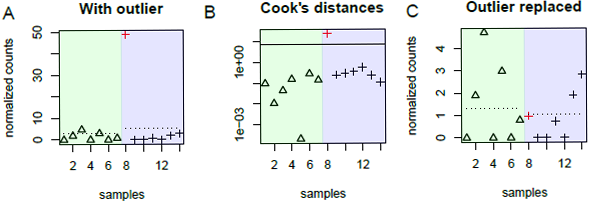
Cook’s distance outlier detection. Shown are normalized counts and Cook’s distances for a 7 by 7 comparison of the Bottomly et al. [15] dataset. (A) Normalized counts for a single gene, samples divided into groups by strains (light green and light blue). Dotted segments represent fitted means. An apparent outlier is highlighted in red. (B) The Cook’s distances for each sample for this gene, and the 99% quantile of the F(p, m – p) cutoff used for flagging outliers. Note the logarithmic scaling of the y-axis. (C) The normalized counts after replacing the outlier with the trimmed mean over all samples, scaled by size factor. The fitted means now are less affected by the single outlier sample.

**Supplementary Figure S5:**
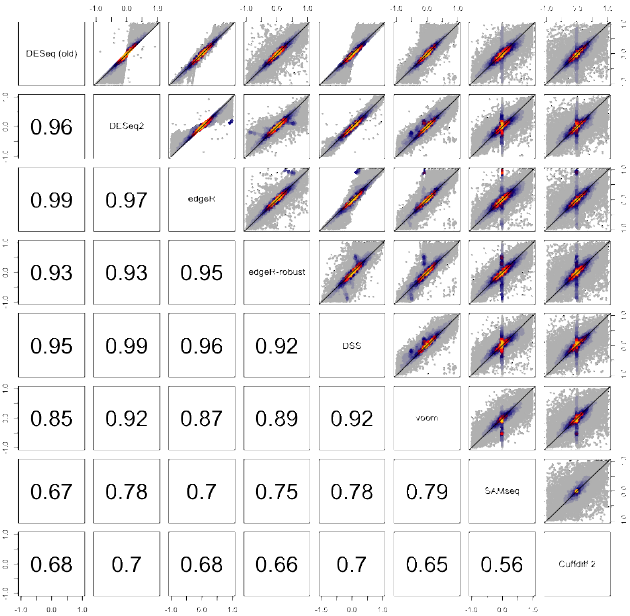
Scatterplots of estimated logarithmic fold changes from all algorithms. log_2_ fold changes are estimated from one of the verification sets of the Bottomly et al. [15] dataset (see section *Benchmarks on RNA-seq data*). Bottom panels display the Pearson correlation coefficients. We note that the direction of the estimate of differential expression for *DESeq2* and *Cuffdiff 2* accorded for the majority of genes called differentially expressed: Among genes which were called differentially expressed by either of these two algorithms, both agreed on the sign of the estimated logarithmic fold change for 96% of genes (averaged over all 30 replicates) in the evaluation set and for 96% of genes in the verification set.

**Supplementary Figure S6:**
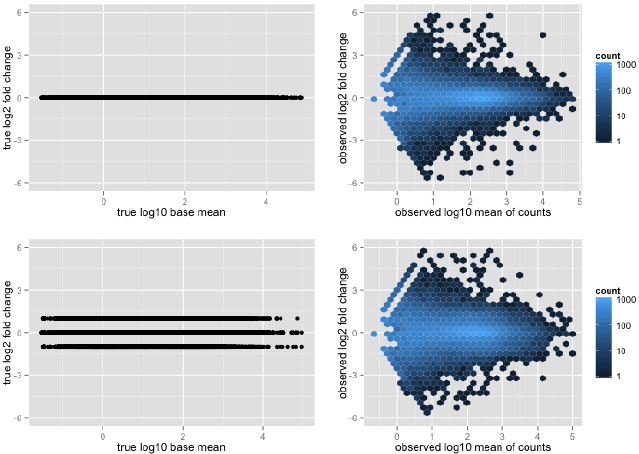
True logarithmic fold changes and the observed logarithmic fold changes induced by the simulation for differential expression. The left plots show the true logarithmic fold changes and true logarithm of base mean, while the right plots show the observed logarithmic fold changes and observed logarithm of the mean of counts for a 4 vs 4 sample comparison. The observed logarithmic fold change was calculated as the logarithm of the mean of counts in one group divided by the mean of counts in the second group. In the top row, all true logarithmic fold changes were equal to zero. On the bottom row, 20% of true logarithmic fold changes were set to a fixed value as in the simulation benchmark for differential expression. We note that mean-independent fixed fold changes produced an MA-plot of observed logarithmic fold changes with mean dependence which is similar to that seen in real data, as in Supplementary Figure S7.

**Supplementary Figure S7:**
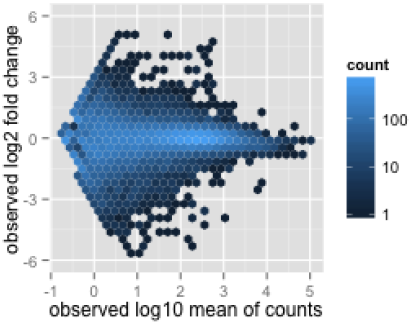
MA-plot from real data. The observed logarithmic fold changes were generated from a 4 vs 4 sample comparison of the Pickrell et al. dataset, wherein there was no known phenotypic difference dividing the groups. The observed logarithmic fold change was calculated as the logarithm of the mean of normalized counts in one group divided by the mean of normalized counts in the second group. The observed mean of counts was calculated as the mean of normalized counts across all samples.

**Supplementary Figure S8:**
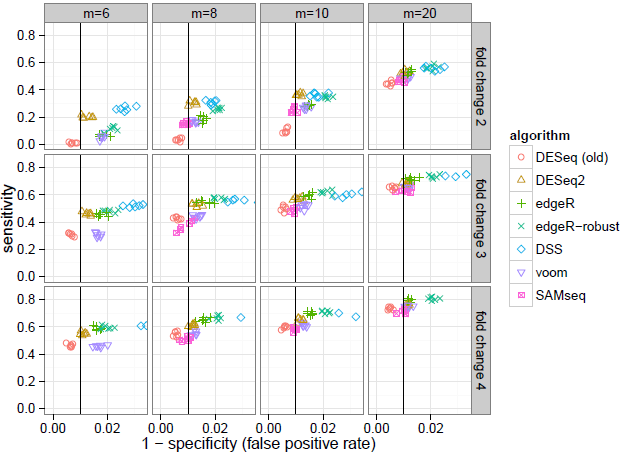
Use of simulation to assess the sensitivity and specificity of algorithms across combinations of sample size and effect size. Shown are results for the benchmark through simulation described in the main text and in Figure 6. The sensitivity was calculated as the fraction of genes with adjusted *p*–value less than 0.1 among the genes with true differences between group means. The specificity was calculated as the fraction of genes with p-value greater than 0.01 among the genes with no true differences between group means. The p-value was chosen instead of the adjusted *p*–value, as this allows for comparison against the expected fraction of p-values less than a critical value given the uniformity of p-values under the null hypothesis. *DESeq2* often had the highest sensitivity of those algorithms which control the false positive rate, i.e., those algorithms which fall on or to the left of the vertical black line (1% *p*–values less than 0.01 for the non-DE genes). *EBSeq* results were not included in this plot as it returns posterior probabilities, which unlike *p*–values are not expected to be uniformly distributed under the null hypothesis.

**Supplementary Figure S9:**
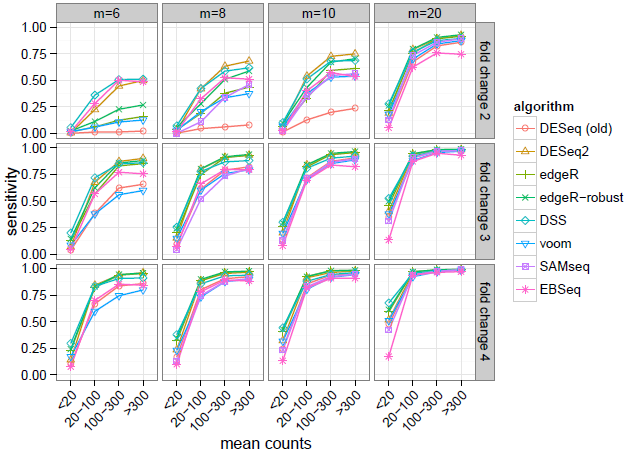
The dependence of sensitivity on the mean of counts for a gene in simulated data. Shown are results for the benchmark through simulation described in Figure 6 and Supplementary Figure S8. The sensitivity of algorithms across combinations of sample size and effect size in the simulated datasets is further stratified by the mean of counts of the differentially expressed genes. The height of the sensitivity curves in this figure corresponds to those shown in Figure 6 and Supplementary Figure S8 which demonstrates the total sensitivity of each algorithm. Points indicate the average over 6 replicates. All algorithms show an expected dependence of sensitivity on the mean of counts. We note that *EBSeq* version 1.4.0 by default removes low count genes – whose 75% quantile of normalized counts is less than 10 – before differential expression calling.

**Supplementary Figure S10:**
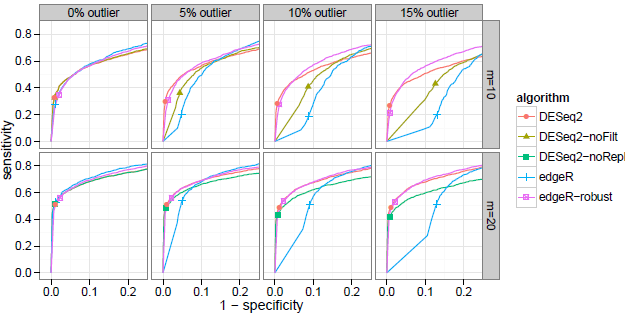
Sensitivity-specificity curves for detecting true differences in the presence of outliers. Negative Binomial counts were simulated for 4000 genes and total sample sizes (m) of 10 and 20, for a two-group comparison. 80% of the simulated genes had no true differential expression, while for 20% of the genes true logarithmic (base 2) fold changes were randomly drawn from {−1, 1}. The number of genes with simulated outliers was increased from 0% to 15%. The outliers were constructed for a gene by multiplying the count o a single sample by 100. Sensitivity and specificity were calculated by thresholding on p-values. Points indicate an adjusted *p*–value cutoff of 0.1. *DESeq2* with the default settings and *edgeR* with the robust setting had higher area under the curve compared to running *edgeR* without the robust option, turning off *DESeq2* gene filtering, and turning off *DESeq2* outlier replacement. *DESeq2* filters gene with potential outliers for samples with 3 to 6 replicates and replaces outliers for samples with 7 or more replicates, hence the filtering can be turned off for the top row (m = 10) and the replacement can be turned off for the bottom row (m = 20).

**Supplementary Figure S11:**
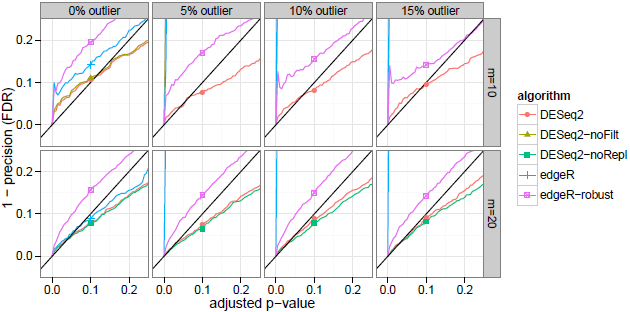
Outlier handling: One minus the precision (false discovery rate) plotted over various thresholds of adjusted *p*–value. Shown are the results for the same simulation with outliers described in Supplementary Figure S10. Points indicate an adjusted *p*–value cutoff of 0.1. *edgeR* run with the robust setting had false discovery rate generally above the nominal value from the adjusted *p*–value threshold (black diagonal line). *DESeq2* run with default settings was generally at or below the line, which indicated control of the false discovery rate.

**Supplementary Figure S12:**
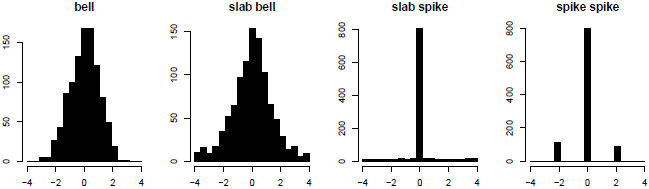
Benchmarking LFC estimation: Models for simulating logarithmic (base 2) fold changes. For the *bell* model, true logarithmic fold changes were drawn from a Normal with mean 0 and variance 1. For the *slab bell* model, true logarithmic fold changes were drawn for 80% of genes from a Normal with mean 0 and variance 1 and for 20% of genes from a Uniform distribution with range from −4 to 4. For the *slab spike* model, true logarithmic fold changes were drawn similarly to the *slab bell* model except the Normal is replaced with a spike of logarithmic fold changes at 0. For the *spike spike* model, true logarithmic fold changes were drawn according to a spike of logarithmic fold changes at 0 (80%) and a spike randomly sampled from −2 or 2 (20%). These spikes represent fold changes of 1/4 and 4, respectively.

**Supplementary Figure S13:**
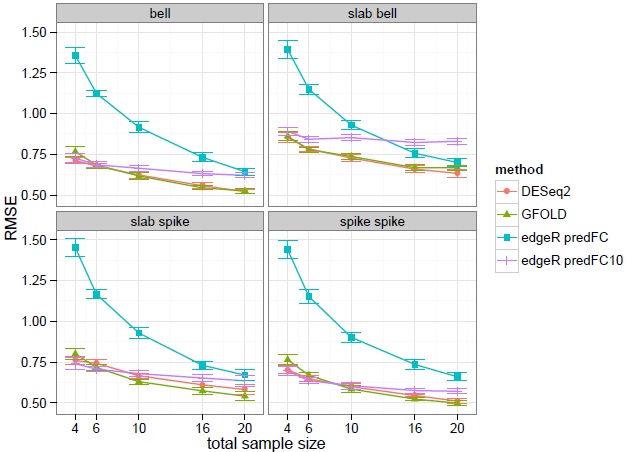
Root mean squared error (RMSE) for estimating logarithmic fold changes under the four models of logarithmic fold changes and varying total sample size *m*. Simulated Negative Binomial counts were generated for two groups and for 1000 genes. Points and error bars are drawn for the mean and 95% confidence interval over 10 replicates. *DESeq2* and *GFOLD*, which both implement posterior logarithmic fold change estimates, had lower root mean squared error to the true logarithmic fold changes over all genes, compared to predictive logarithmic fold changes from edgeR, either using the default value of 0.125 for the *edgeR* argument *prior.count*, or after increasing *prior.count* to 10 *(edgeR* predFC10).

**Supplementary Figure S14:**
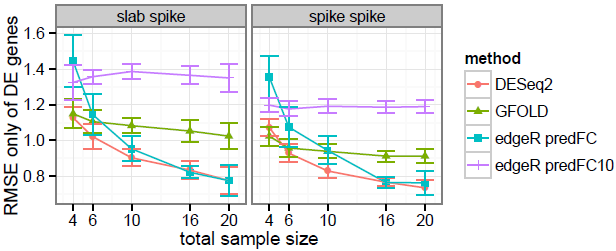
Root mean squared error (RMSE) of logarithmic fold change estimates, only considering genes with non-zero true logarithmic fold change. For the same simulation as shown in Supplementary Figure S13, shown here is the error only for the 20% of genes with non-zero true logarithmic fold changes (for *bell* and *slab bell* all genes have non-zero logarithmic fold change). *DESeq2* had generally lower root mean squared error, compared to *GFOLD* which had higher error for large sample size and to *edgeR* which had higher error for low sample size.

**Supplementary Figure S15:**
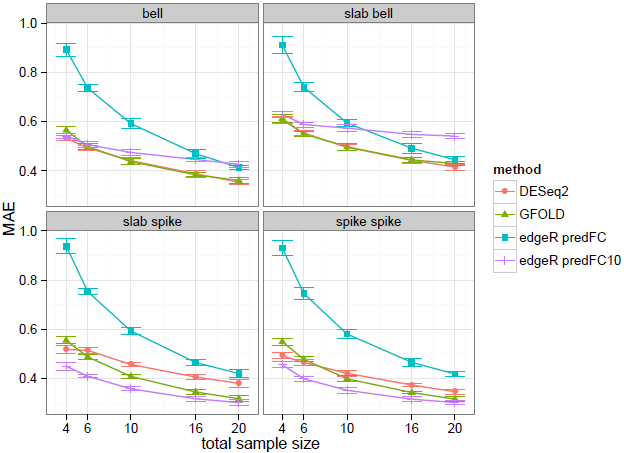
Mean absolute error (MAE) of logarithmic fold change estimates. Results for the same simulation as shown in Supplementary Figure S13, however here using mean absolute error in place of root mean squared error. Mean absolute error places less weight on the largest errors. For the *bell* and *slab bell* models, *DESeq2* and *GFOLD* had the lowest mean absolute error, while for the *slab spike* and *spike spike* models, *GFOLD* and *edgeR* with a *prior.count* of 10 had lowest mean absolute error.

**Supplementary Figure S16:**
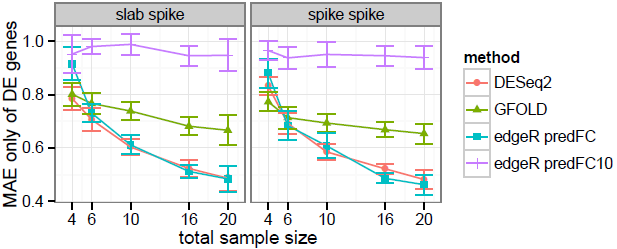
Mean absolute error (MAE) of logarithmic fold change estimates, only considering those genes with non-zero true logarithmic fold change. While in Supplementary Figure S15, considering all genes for the *slab spike* and *spike spike* models, *GFOLD* and *edgeR* with a *prior.count* of 10 had lowest mean absolute error, the mean absolute error for these methods was relatively large for large sample size, when considering only the 20% of genes with true differentially expression. *DESeq2* and *edgeR* generally had the lowest mean absolute error.

**Supplementary Figure S17:**
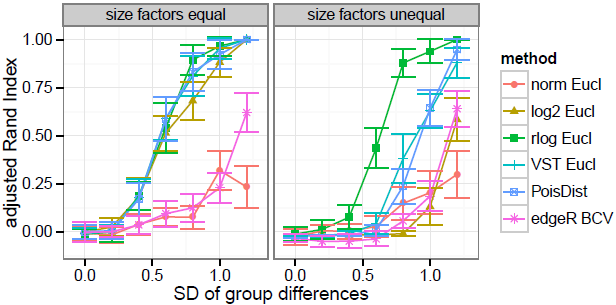
Adjusted Rand Index of clusters using various transformation and distances compared to the true clusters from simulation. 4 simulated clusters with 4 samples each were generated using Negative Binomial counts over 2000 genes using the means and gene-wise estimates of dispersion from the Pickrell et al. dataset. 80% of genes were given equal mean across clusters, while for 20% of genes, logarithm (base 2) fold changes from a centroid were drawn from a zero-centered Normal distribution while varying the standard deviation (SD, x-axis). Larger standard deviation resulted in more distinct clusters, which are easier for the methods to recover. Simulation was performed with equal size factors, and with size factors for each group set to [1,1,1,3]. The methods assessed were: Euclidean distance on counts normalized by size factor, logarithm of normalized counts plus a pseudocount of 1, rlog transformed counts and variance stabilized counts (VST). Additionally, the Poisson Distance from the *PoiClaClu* package and the Biological Coefficient of Variation (BCV) distance from the *plotMDS* function of the *edgeR* package were used for hierarchical clustering. We note that the default distance used by *plotMDS* is not the BCV distance but more similar to the Euclidean distance of logarithmic counts. The points and error bars indicate the mean and 95% confidence interval from 20 replicates. In the simulations with equal size factors, the Poisson distance, the VST and the rlog had the highest accuracy in recovering true clusters. In the unequal size factor simulations, the rlog outperformed the other methods.

**Supplementary Figure S18:**
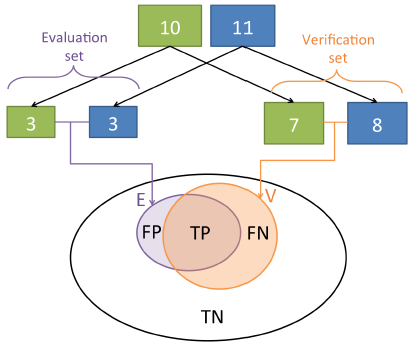
Diagram of the evaluation scheme for the benchmarks using real RNA-seq data. The Bottomly et al. dataset with 10 and 11 replicates was split into a 3 vs 3 “evaluation set” and a 7 vs 8 “verification set”. The positive calls from the verification set, denoted as set V, were taken as a pseudo-gold standard of truly differentially expressed genes. The algorithms were then evaluated based on the set E of positive calls in the evaluation set, comparing to the gold-standard calls from the set *V*. Sensitivity was calculated as |*E* ∩ *V*|/|*V*| and precision was calculated as |*E* ∩ *V*|/|*E*|. Each algorithm’s calls in the evaluation set were compared against each algorithm’s calls in the verification set.

**Supplementary Figure S19:**
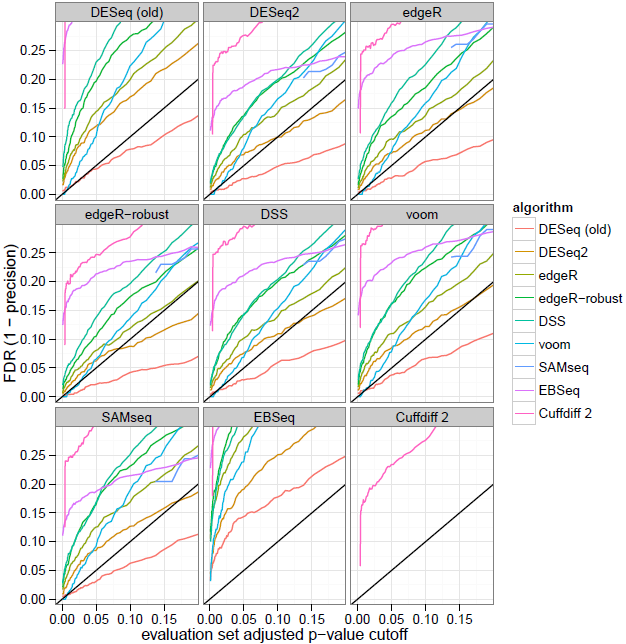
Actual versus nominal false discovery rate for the Bottomly et al. dataset. The actual false discovery rate was calculated using the median of (1 – precision), though here varying the adjusted *p*–value cutoff, i.e., the nominal FDR, for the evaluation set (for *EBSeq*, the posterior probability of equal expression was used). A false positive was defined as a call in the evaluation set for a given critical value of adjusted *p*–value which did not have adjusted *p*–value less than 0.1 in the verification set. Ideally, curves should fall on the identity line (indicated by a black line); curves that fall above indicate that an algorithm is too permissive (anti-conservative), curves falling below indicate that an algorithm does not use its type-I error budget, i.e., is conservative. *DESeq2* had a false discovery rate nearly matching the nominal false discovery rate (black diagonal line) for the majority of algorithms used to determine the verification set calls. The old *DESeq* tool was often too conservative.

**Supplementary Figure S20:**
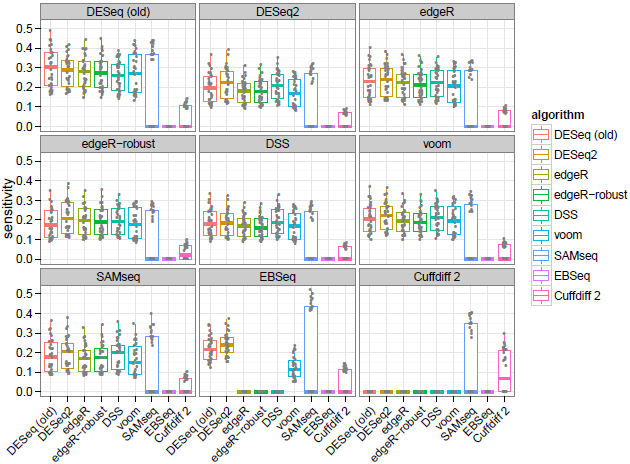
Sensitivity of algorithms evaluated while controlling the median precision. While it was generally noted that sensitivity and precision were negatively correlated (Figure 8 and Figure 9), here this effect was controlled by setting the adjusted *p*–value cutoff for the evaluation set calls such that the median precision of all algorithms would be 0.9 (actual false discovery rate of 0.1). This amounted to finding the point on the x-axis in Supplementary Figure S19, where the curve crosses 0.1 on the y-axis. For most algorithms, this meant setting an adjusted *p*–value cutoff below 0.1. *DESeq2* often had the highest median sensitivity for a given target precision, though the variability across random replicates was generally larger than the difference between algorithms.

**Supplementary Figure S21:**
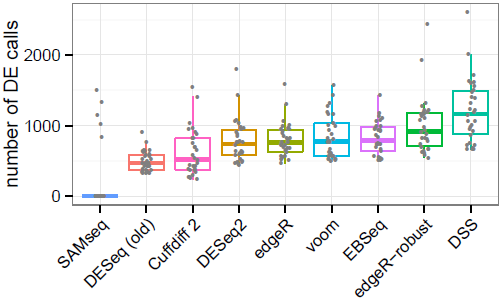
**Number of total calls in the evaluation set (3 vs 3 samples)** of the sensitivity/precision analysis using the Bottomly et al. [15] dataset thresholding at adjusted *p*–value < 0.1, over 30 replications.

**Supplementary Figure S22:**
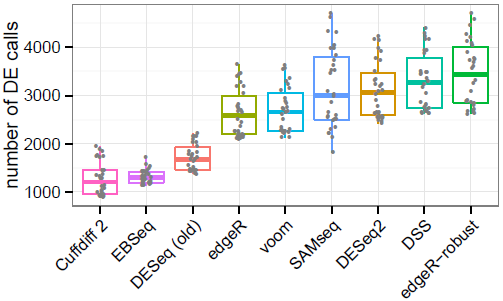
**Number of total calls in the verification set (7 vs 8 samples)** of the sensitivity/precision analysis using the Bottomly et al. [15] dataset thresholding at adjusted *p*–value < 0.1, over 30 replications.

**Supplementary Figure S23:**
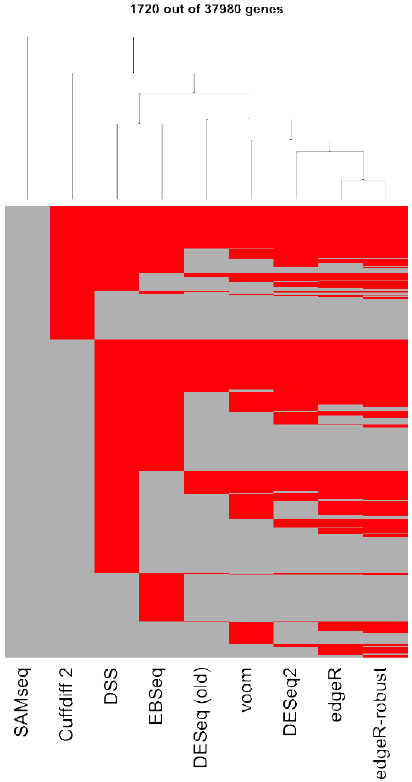
Clustering of each algorithm’s calls on the evaluation set (3 vs 3 samples) for one replicate of the sensitivity/precision benchmark. Genes are on the vertical axis and algorithms on the horizontal axis. Red lines indicate a gene had adjusted *p*–value < 0.1 in the evaluation set. Genes in which no algorithm had a call are not shown. Clustering is based on the Jaccard index.

**Supplementary Figure S24:**
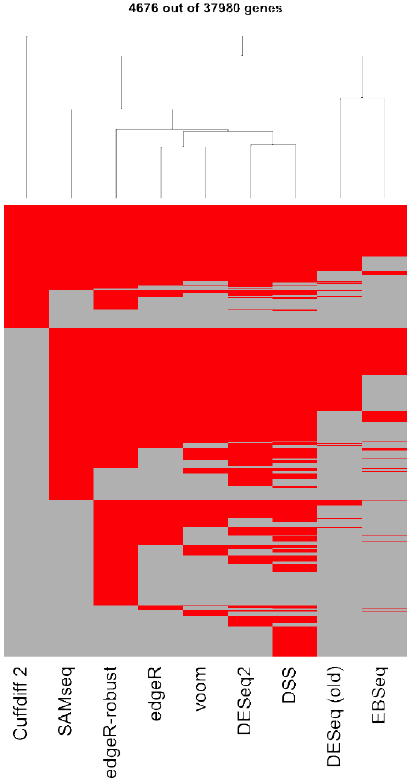
Clustering of algorithm calls on the verification set (7 vs 8 samples) for one replicate of the sensitivity/precision benchmark. Genes are on the vertical axis and algorithms on the horizontal axis. Red lines indicate a gene had adjusted *p*–value <0.1 in the verification set. Genes in which no algorithm had a call are not shown. Clustering is based on the Jaccard index.

**Supplementary Figure S25:**
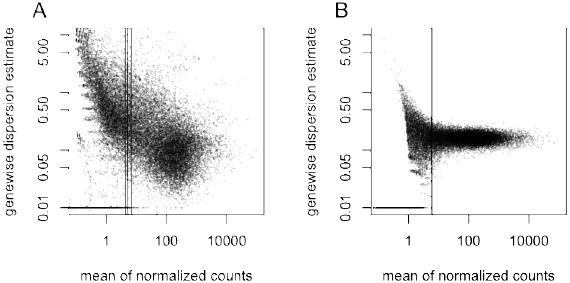
Demonstration through simulation that the dependence of dispersions on the mean seen in Figure 1B is not an artifact of estimation bias. (A) The gene-wise estimates of dispersion for the 69 samples of the Pickrell et al. dataset. (B) The gene-wise estimates of dispersion for a simulated Negative Binomial dataset, using a fixed dispersion of *α* = 0.16, equal to the asymptotic gene-wise dispersion estimate *α_0_* seen in the original dataset (A), and with the same means and the same number of genes and samples as the original dataset. Genes with dispersion estimates below the plotting range are depicted at the bottom of the frame. For genes with mean counts greater than ∼5, the gene-wise dispersion estimates do not exhibit a dependence on the mean count for the simulated data in panel B. Vertical lines indicate the reciprocal of the asymptotic dispersion ao, on the scale of raw counts for the 1^st^, 2^nd^ and 3^rd^ quartile of the size factors.

**Supplementary Figure S26:**
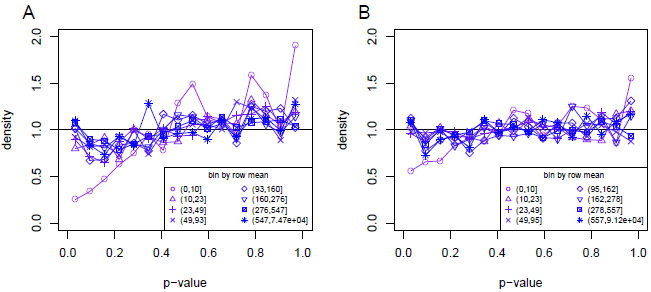
Marginal null histogram of the test statistic, *p*–values, conditioning on the filter statistic, the row mean of normalized counts across all samples, used for independent filtering. A simulated dataset was constructed with (A) 6 samples or (B) 12 samples and 20,000 genes. In either case the samples were equally divided into 2 groups with no true difference between the means of the two groups. The means and dispersions of the Negative Binomial simulated data were drawn from the estimates from the Pickrell et al. dataset, and the standard *DESeq2* pipeline was run. The histogram of *p*–values was estimated at 16 equally spaced intervals spanning [0,1]. The marginal distributions of the test statistic were generally uniform while conditioning on bins based on the filter statistic. The row mean bin with the smallest mean of normalized counts (mean count 0 – 10) was depleted of small *p*–values. The black line indicates the expected frequency for a Uniform distribution.

